# Tasting the differences: microbiota analysis of different insect-based novel food

**DOI:** 10.1101/2020.02.20.957845

**Authors:** Jessica Frigerio, Giulia Agostinetto, Andrea Galimberti, Fabrizio De Mattia, Massimo Labra, Antonia Bruno

**Author notes:** These authors contributed equally to this work (Jessica Frigerio, Giulia Agostinetto).

## Abstract

Traceability, quality and safety of edible insects are important both for the producers and the consumers. Today, alongside the burst of edible insects in western countries, we are facing a gap of knowledge of insect microbiota associated with the microbial ecosystems of insect-based products. Recent studies suggest that the insect microbiota can vary between insect species and that can be shaped by additional factors, such as rearing conditions. Also, the production processes of raw materials (i.e. insect flour) into final food products can affect the insect microbiota too. This has consequences for the evaluation of food safety and food traceability. In this context, High-Throughput DNA Sequencing (HTS) techniques can give insight into the carryover of insect microbiota into final food products. In this study, we investigated the microbiota composition of insect-based commercial food products, applying HTS techniques coupled with bioinformatic analysis. The aim of this work was to analyse the microbiota variability of different categories of insect-based products made of *A. domesticus* (house cricket), *T. molitor* (mealworm beetle), and *A. diaperinus* (lesser mealworm or litter beetle), including commercial raw materials and processed food items, purchased via e-commerce from different companies. Our data revealed that samples cluster per insect species based on microbiota profile and preliminary results suggested that a small number of prevalent bacteria formed a “core microbiota” characterizing the products depending on the insect, suggesting that a resident microbiota is conserved. This microbial signature can be recognized despite the different food processing levels, rearing conditions selling companies. We showed that differences exist when comparing raw vs processed food made of the same insect, or similar products produced by different companies as well, laying the groundwork for further analyses. These results support the application of HTS analysis for studying the composition of processed insect food in a wider perspective, for food traceability and food quality control.

## 1. Introduction

Entomophagy is an emerging and fashionable diet issue in western countries. Insects are an important source of energy for human diets, thanks to their richness in essential nutrients (Rumpold & Schlüter, 2015). They have a protein content average value ranging from 30% to 65% of the total dry matter, but they are also rich in micronutrients such as iron, zinc and calcium (Dobermann, Swift & Field, 2017). Moreover, preliminary studies of Oonincx & de Boer (Oonincx & de Boer, 2012) stated that, compared to other livestock animals, insect farming has a lower environmental footprint.

Safety, traceability and quality of edible insects are of great interest both for the producers and the consumers, heavily affecting the acceptance of edible insects in the human diet (House, 2016). New tools for quality and safety controls on these food items could also benefit institutions like food agencies, customs and health departments in the evaluation of new product development based on processed insects. Geographically, there are three legal categories. Concerning the Anglo-Saxon countries such as the UK, USA, Canada, New Zealand and Australia, edible insects do not represent a novel food and the local food agencies have authorized their import and sales. In the European Union, the regulation (Regulation EU 2015/2283) has classified edible insects as novel foods, which follow specific rules and require specific authorizations before allowing them to be distributed (Klunder HC, Wolkers-Rooijackers, Korpela, Nout, 2016; Van Huis, 2012; Shutler et al., 2017). Finally, in the remaining areas such as Asian countries, insects are considered traditional food, therefore they are commonly commercialized and consumed. This system of multiple rules could lead to difficulties in the trading of these products. In addition, food safety authorities and the scientific community are discussing whether edible insects can be a reliable solution or a problem to the food security (Belluco et al., 2015, Di Mattia, Battista, Sacchetti, Serafini, 2019).

The potential safety risks of edible insects are chemical hazards including pesticides, heavy metals, allergens, toxins (mycotoxin and bacterial toxins). There is a risk that harmful insect microbes are transmitted through the consumption of insect products (van der Spiegel, Noordam, van der Fels-Klerx, 2013). Most of the insect microbiota are associated with gut (e.g., the intrinsic insect symbionts in the intestinal tract and in the proximity of other anatomical compartments) or related to extrinsic sources, such as environment and rearing conditions (substrates and feed), handling, processing and preservation (ANSES, 2014). Especially, as stressed recently by the European Food Safety Authority (EFSA, 2015), spore-forming bacteria in processed edible insects (including freeze-dried, boiled and dried varieties) can be considered a dangerous source of biological contamination as well.

Today, there is little information available about insect microbiota associated with insect-based products which potentially harbors organisms harmful to human health. Garofalo and colleagues (Garofalo et al., 2017) explored the microbiota of marketed processed edible insects using culture-based methods and pyrosequencing. They described, among others, the microbiota of whole dried small crickets (*Acheta domesticus*) and whole dried mealworm larvae (*Tenebrio molitor*), revealing a great bacterial diversity and variability among individual insect species: some of the identified microbes may act as opportunistic pathogens in humans, such as Listeria spp., Staphylococcus spp., Clostridium spp. and Bacillus spp., while others represent food spoilage bacteria, as well as *Spiroplasma* spp. in mealworm larvae. The insect diet and social behavior have a great impact on the composition of the gut microbial community (Tinker & Ottesen, 2013), therefore different insect farm conditions result in different microbiological ecosystems. Although some authors such as Stoops and co-workers (Stoops et al., 2017) suggested that the microbial taxonomic composition varies mainly with insect species, the additional factors such as the growing substrates or contact with soil may play an important role in the composition of the insect gut microbiota (Klunder HC, Wolkers-Rooijackers, Korpela, Nout, 2016; EFSA, 2015; Li et al., 2016). Considering the insect production system, industrial practices, such as post-harvest starvation and rinsing, can affect the microbial quality of the final insect products too (Wynants et al., 2018). Since all food products, including those insect-based, undergo processing, the risk for human safety should be measured throughout the various stages, from raw materials (i.e. insect flour) to final food products. High-Throughput DNA Sequencing (HTS) offers a standardized and sensitive method to evaluate the microbial community changes by analysing a wide range of food products (De Filippis, Parente & Ercolini, 2019). The search for a microbial signature represents both an opportunity to verify food safety and food traceability strategy, indeed the microbial variation gives insight about rearing and processing products. The microbial variability allows to obtain more information besides the identification of the insect species identification, like the hygienic and sanitary conditions concerning the rearing systems. Moreover, the insect microbiota can be used to identify the geographical origin of a food product and used as a tracing signature, as previously demonstrated by Mezzasalma and colleagues (Mezzasalma et al., 2017).

In this study, we evaluated the microbiota composition of insect-based commercial food products, applying HTS with complementary bioinformatics analysis. The aim of this study was to analyse the microbiota variability of different categories of insect-based products (including commercial raw materials and processed food items), purchased via e-commerce from different companies. We sought to define with a preliminary study if HTS-based tools could be useful to get insight into the impact of the food processing steps on the transfer of the insect microbiota into the final product.

## 2. Materials and methods

### 2.1 Insect food products

A total of 12 commercial insect-based products were purchased via e-commerce from six different companies (Table 1). Referring to the label information, these products contained only one insect species each, belonging to the orders Orthoptera (*Acheta domesticus*), and Coleoptera (*Alphitobius diaperinus* and *Tenebrio molitor*) (S1 Table).

**Table 1.**
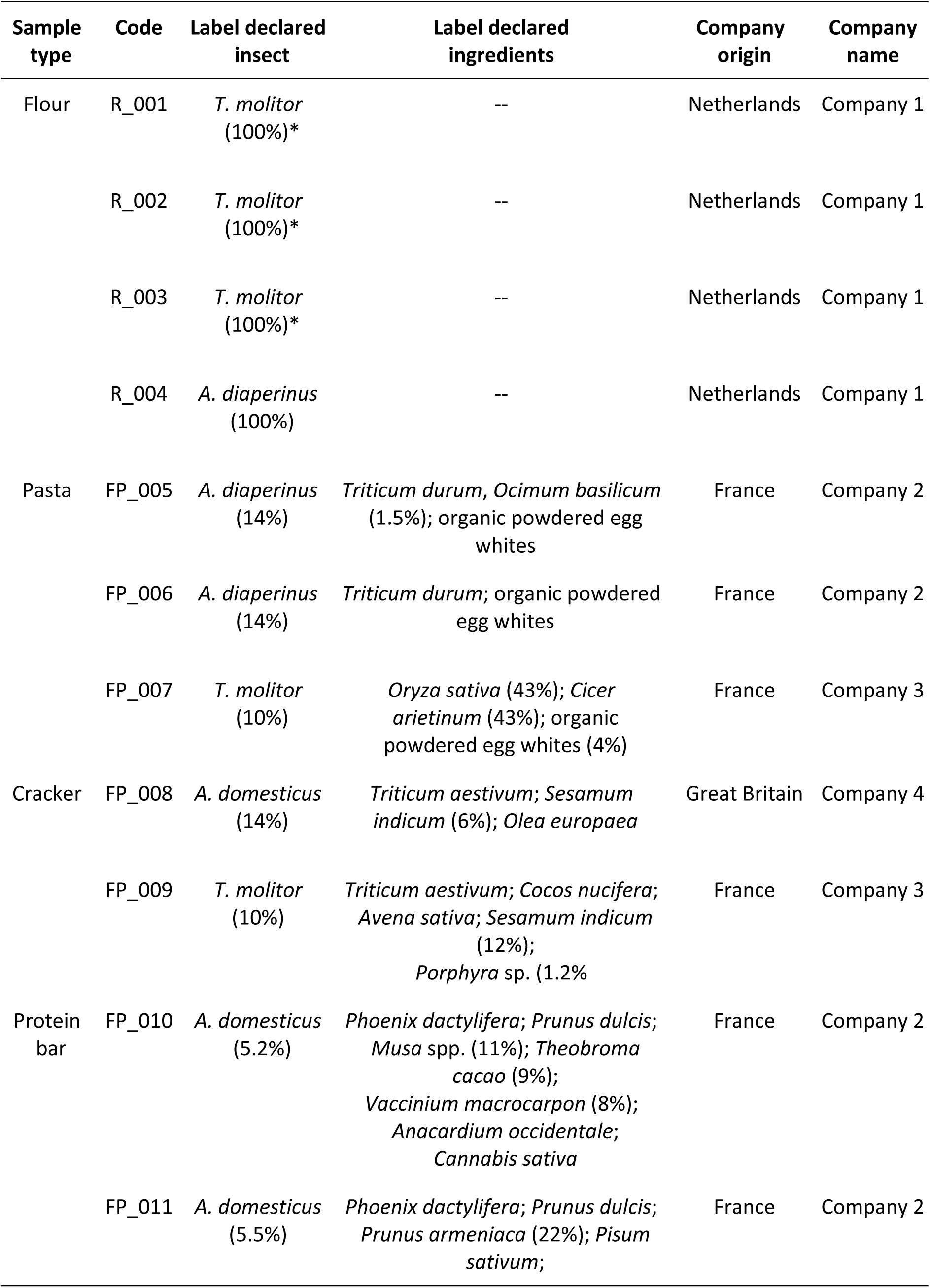
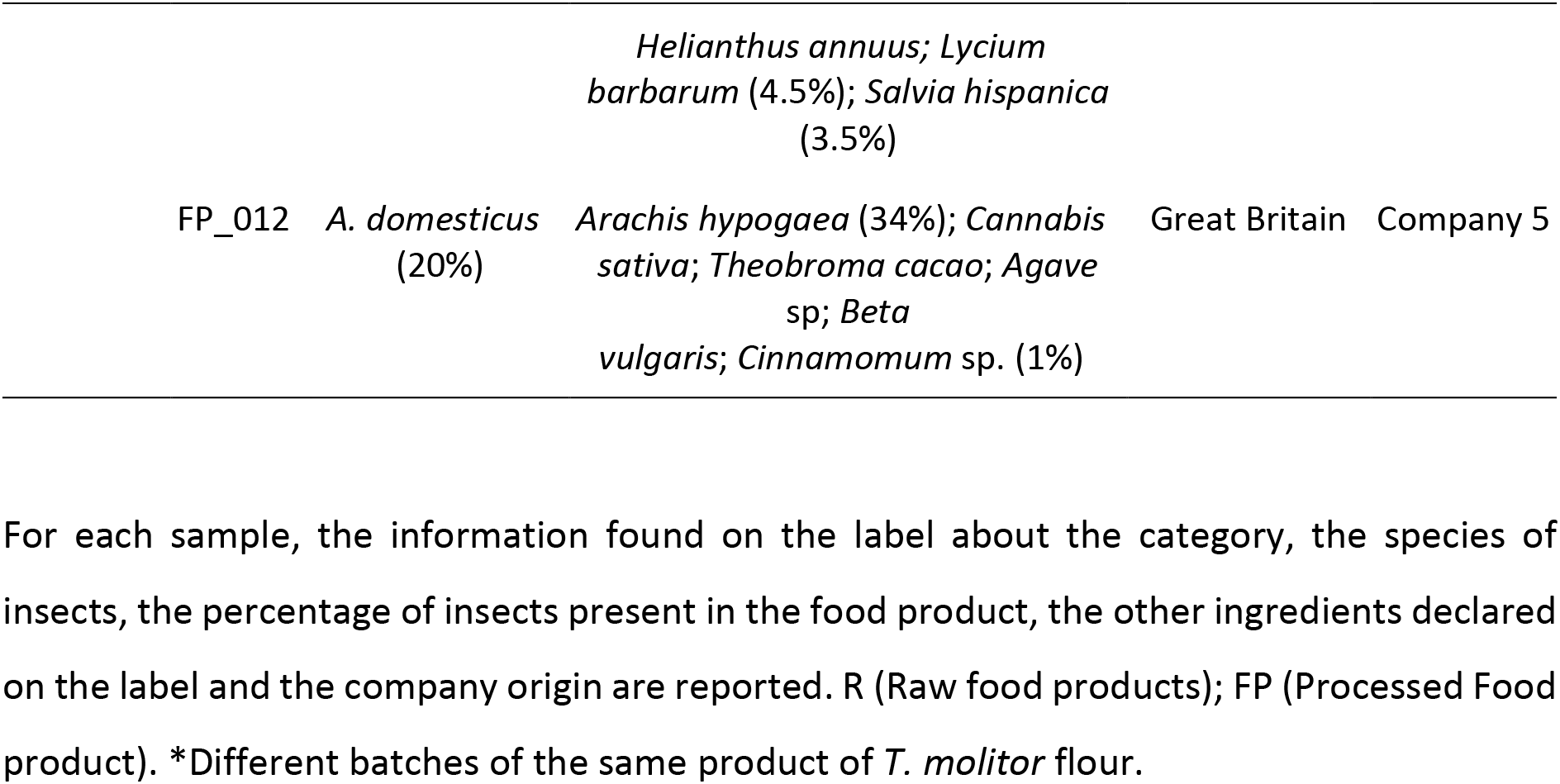
List of analysed insect-based products.

Among these, four products were pure insect flours belonging to the species *Alphitobius diaperinus* (n=1) and *Tenebrio molitor* (n=3), and they have been categorized as insect raw material. In the case of *T. molitor*, we collected three different batches of the same product to test if any variability among batches exists, considering microbiota composition (samples R_001; R_002 and R_003). The remaining eight samples represented processed food products: pasta (n=3), crackers (n=2) and protein bars (n=3).

### 2.2 DNA extraction

High-quality genomic DNA was obtained starting from 250 mg of each sample of table 1 using DNeasy PowerSoil Kit (QIAGEN, Hilden, Germany), according to manufacturer’s instructions. Three replicates of DNA extraction were generated for each sample plus a negative control. Purified DNA was checked for concentration and purity by using a Qubit 2.0 Fluorometer and Qubit dsDNA HS Assay Kit (Invitrogen, Carlsbad, California, United States).

### 2.3 DNA barcoding characterization of insect samples

The 658 bp mtDNA COI region was used to validate the animal species declared on the label in the sampled insect-based products. This region was amplified and sequenced for all 12 samples according to the primer pairs presented by Folmer and colleagues (Folmer et al., 1994) and the protocol described in Bellati et al. (Bellati et al., 2014). Each sequence was defined the nearest matches with the BLAST algorithm using the following cut-off values/maximum identity >99% and query coverage of 100%.

### 2.4 HTS library preparation and sequencing

To characterize the bacterial composition of the investigated insect-based products, 16S rRNA genes (V3 and V4 hypervariable regions) of the obtained gDNA extracts were sequenced using a High-Throughput Sequencing approach. Amplicons were generated following the protocol described by Caporaso et al. (Caporaso et al., 2012) with Illumina adapters (S2 Table), with minor modifications as described in Frigerio et al. (Frigerio et al., 2020): we used puReTaq Ready-To-Go PCR beads (GE Healthcare Life Sciences, Italy) according to manufacturer’s instructions in a 25 μL reaction, containing 1 μL 10 mM of each primer and up to 50 ng of gDNA. The amplification profile consisted of an initial denaturation step for 5 min at 95 °C, followed by 25 cycles of denaturation (30 s at 95 °C), annealing (30 s at 55 °C), and elongation (30 s at 72 °C), and finally elongation at 72 °C for 5 min. Amplicon DNA was checked for concentration by using a Qubit 2.0 Fluorometer and Qubit dsDNA HS Assay Kit (Invitrogen, Carlsbad, California, United States) and amplicon length was measured by comparison against QX DNA Size Marker using the Qiaxcel Automatic electrophoresis system (QIAGEN, Hilden, Germany). Samples were sequenced by the Center for Translational Genomics and BioInformatics (Milan, Italy). The sequencing was performed on the MiSeq sequencing platform (Illumina, San Diego, CA, USA) with a paired-end approach (MiSeq Reagent Kit v3, 2 x 300 bp).

### 2.5 Bioinformatics analysis

Illumina reads were analysed with QIIME2, Quantitative Insights Into Microbial Ecology 2 program (ver. 2019.4; https://qiime2.org/) (Boylen et al., 2018). Sequences were demultiplexed with native plugin and DADA2 (Divisive Amplicon Denoising Algorithm 2) (Callahan et al., 2016) was applied to obtain ASVs sequences (or features) (Callahan et al., 2017), trimming primers and performing a quality filter with an expected error of 2.0. Chimeric sequences were removed using the consensus method. Features with at least 10 representatives associated and detected in at least two samples were kept. The taxonomic assignment of representative sequences was carried out using the *feature-classifier* (https://github.com/qiime2/q2-feature-classifier) plugin implemented in QIIME2, using *classify-consensus-vsearch* method against the SILVA SSU non-redundant database (132 release), adopting a consensus confidence threshold of 0.8. Taxa bar plots were generated with the QIIME2 dedicated plugin *taxa* (https://github.com/qiime2/q2-taxa). As ASVs assigned to Cyanobacteria phylum (class Chloroplast) were considered potential plant contaminants, they were removed from the downstream analysis. Reads of mitochondrial or eukaryotic origin were also excluded. Overlap among technical replicates was calculated considering taxa at family level weighted for abundances (Wen et al., 2017). Alpha diversity was carried out considering presence/absence of ASVs and Shannon index. Statistical differences among samples belonging to the same insect species were calculated using alpha-group-significance plugin by QIIME2, performing also a pairwise contrast (Kruskal and Wallis, 1952). Beta diversity, instead, was carried out considering qualitative (Jaccard and unweighted UniFrac) and quantitative (Bray-Curtis and weighted UniFrac) distance metrics (Lozupone et al., 2011), using QIIME2 *core-metrics* plugin (https://github.com/qiime2/q2-diversity). Statistical differences were calculated by permutation-based ANOVA (PerMANOVA) functions of *beta-group-significance* plugin (Anderson, 2001), with 999 permutations, considering insect species and sample type categories. A PerMANOVA Pairwise contrast was performed with *beta-group-significance* plugin. Principal coordinates plots (PCoA) method was used to explore the structure of microbial communities. The phylogenetic tree necessary to calculate UniFrac distances was built on the alignment of ASVs representative sequences using align-to-tree-mafft-fasttree method by *phylogeny* plugin (https://github.com/qiime2/q2-phylogeny). Heatmap visualization was used to explore the abundance of bacteria families among samples and was generated by QIIME2. Core microbiota among insect samples was calculated considering the ceiling of the mean of species frequencies among samples and keeping a core threshold of 0.7 (minimum fraction of samples that a species must be observed in), performed with *core-features* plugin (https://github.com/qiime2/q2-feature-table). A Venn diagram was created starting from core microbiota results setting the threshold = 1, by calculating the number of shared and unique taxa per insect collapsed at the genus level.

## 3. Results

### 3.1 Sequencing output

All the replicates of the 12 collected samples showed good DNA quality (i.e., A260/A230 and A260/A280 absorbance ratios within the range 1.6 - 2.2) and good yield (20-40 ng/μl). The DNA barcoding (mt COI) sequencing results indicated that all the tested samples were composed of insects. Moreover, the BLAST analysis against reference insect DNA barcoding sequences confirmed that all samples corresponded with the declared insect species (i.e., maximum identity > 99% with the declared species).

HTS analysis produced about 8,573,268 raw reads from the analysed samples, with an average of 231,709 reads per sample (DS = 128,393). After quality filtering, merging reads, chimera and contaminants removal, we obtained a total of 605 ASVs (Amplicon Sequence Variants) [26]. Negative controls (deriving from DNA extraction and amplification step) for library sequencing were not included in the analysis since they encompassed a very low number of DNA reads.

### 3.2 Microbial diversity analysis

From overlap calculations for technical replicates, family overlap resulted in a mean of 96%, with a standard deviation of 0.06.

Both considering ASVs and Shannon metric, differences among samples derived from different insects were observed (Table S3).

Samples belonging to raw material (flour) and food products (crackers, pasta and protein bars) showed a significant difference, considering both qualitative (Jaccard and Unweighted UniFrac) and quantitative metrics (Bray-Curtis and Weighted UniFrac) (p-value<0.01). Overall, we observed a significant difference among samples belonging to different insects (p-value<0.01; q-value<0.01).

### 3.3 Taxonomic composition analysis

A total of 9 bacterial phyla, 14 classes, 34 orders, and 67 families were identified (Fig 1, S5 Table).

**Fig 1.**
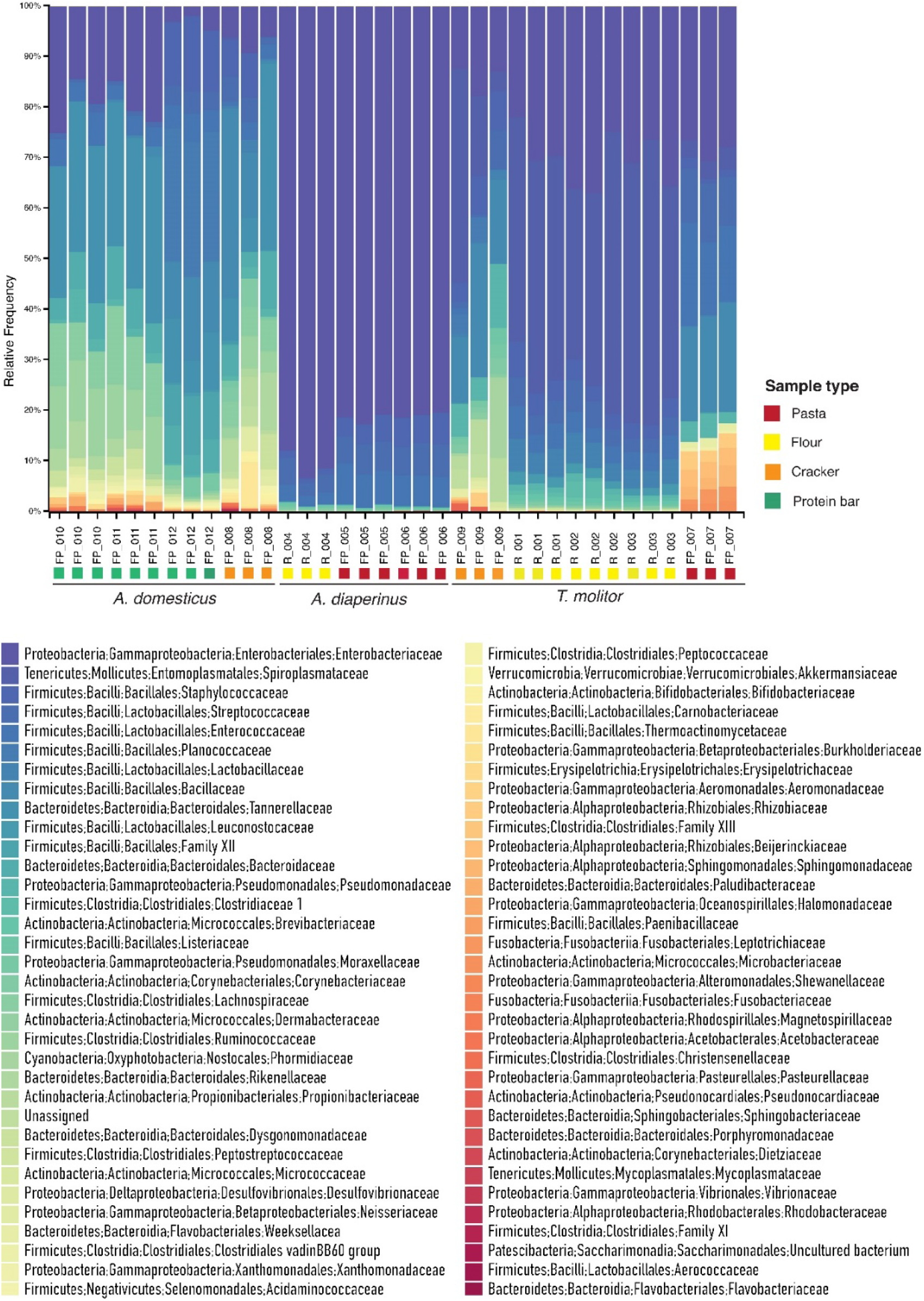
Relative abundance of bacteria families recovered in the insect-based products through 16S metabarcoding sequencing. Bacteria families are reported in gradient colours indicating relative abundances. For each sample, the sample type is reported (pasta: red square; flour: yellow square; cracker: orange square; protein bar: green square).

Taxonomic analysis revealed that most of the sequences in all the samples were associated with the phyla Proteobacteria (47%) and Tenericutes (26%), followed by Firmicutes (23%). 13% resulted in Unassigned taxa. Looking inside the taxonomic rank of class, the most abundant were Gammaproteobacteria, with 47% of sequences, Mollicutes (26%), and Bacilli (22%). Enterobacteriales was the most abundant order, encompassing 45% of the sequences, distributed across all the samples, followed by Entomoplasmatales (26%), Lactobacillales (12%), Bacillales (10%), and Bacteroidales (2.6%). On the whole, the remaining 29 orders covered 4.4% of sequences. The Enterobacteriaceae family accounted for 45% of sequences, whereas Spiroplasmataceae represented 26% of sequences.

Considering taxa distribution per insect (Fig 2), we can notice differences in microbial composition, spanning from the phylum level to a deeper taxonomic resolution. Considering taxonomy per insect species, at the taxonomic level of order, we found that *A. domesticus*-based samples were dominated by Bacillales (54%), followed by Bacteroidales (21.2%), and Lactobacillales (8.9%), representing 84.1% of 28 orders. On the other hand, food products made with *A. diaperinus* had most of the sequences assigned to Enterobacteriales (89.6%), with the remaining 7% and 2.1% assigned to Lactobacillales and Bacillales, respectively, and 1.3% of sequences distributed in 11 orders. *T. molitor*-based food products showed 45.4% of sequences corresponding to Entomoplasmatales order, 29.5% to Enterobacteriales, 14.5% to Lactobacillales, and the remaining 10.6% to 21 different orders.

**Fig 2.**
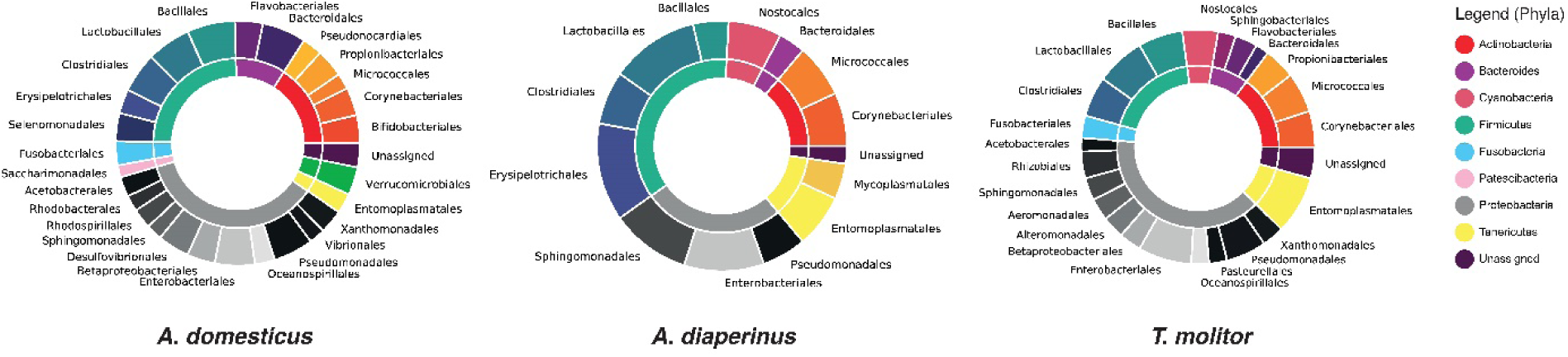
Donuts charts of *A. domesticus, A. diaperinus*, and *T. molitor* microbial composition. Phyla in the inner circle and Orders in the outer circle are reported. Abundances are expressed as log frequency, in order to better show underrepresented taxa.

Focusing on specific features, we observed that the most abundant feature was assigned to an uncultured Spiroplasma (25%), reported exclusively in *T. molitor* samples. The fifth most abundant feature (3%), assigned to the genus *Kurthia* (Planococcaceae; Bacillales; Bacilli; Firmicutes) was detected only in *A. domesticus* protein bars produced by the British company 5, but not in samples belonging to the British company 4. Moreover, all and only the food products deriving from British company 5 are characterized by the presence of a specific feature assigned to *Exiguobacterium* (Family XII; Bacillales; Bacilli; Firmicutes). Considering features shared between protein bars belonging to British company 5 and French company 2, the most abundants were assigned to Enterobacteriaceae family (20,4%) (Enterobacteriales; Gammaproteobacteria; Proteobacteria), followed by Tannerellaceae (14,3%) (Bacteroidales; Bacteroidia; Bacteroidetes) and Lachnospiraceae (14,3%) (Clostridiales; Clostridia; Firmicutes).

A feature assigned to an uncultured *Parabacteroides* (Tannerellaceae; Bacteroidales; Bacteroidia; Bacteroidetes) is unique for *A. domesticus* samples, whereas features assigned to *Enterobacter* (Enterobacteriaceae; Enterobacteriales; Gammaproteobacteria; Proteobacteria), a different microorganisms belonging to Enterobacteriaceae, and *Enterococcus* (Enterococcaceae; Lactobacillales; Bacilli; Firmicutes) were found only in *A. diaperinus* food products.

To better visualize the microbial variation among different food products, and which family mostly contribute to distinguish food products, a heatmap based on relative abundances was generated (Fig 3).

**Fig 3.**
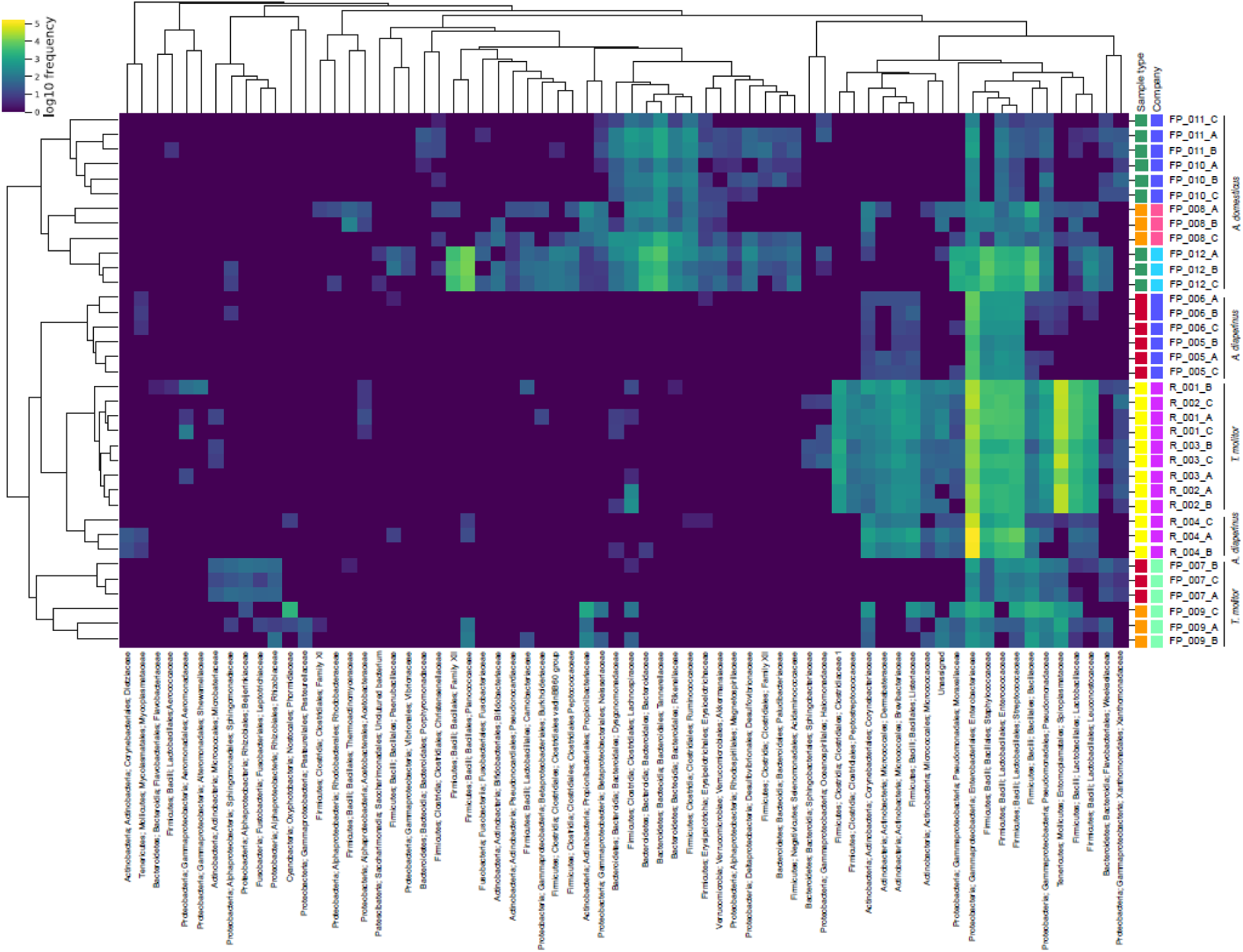
Heat map diagram showing the relative abundance of families for each sample. Colour shading in the heat map indicates the abundance, expressed as log10 frequency, of each family in the sample. Samples type category are flour (yellow), pasta (red), cracker (orange) and protein bar (green). Companies are represented in fuchsia (Company 1), blue (Company 2), aquamarine (Company 3), pink (Company 4) and light blue (Company 5).

Analysing the sample cluster dendrogram, two main clusters separate samples based on insect order, composed by *A. domesticus* (Orthoptera) food products and *T. molitor* plus *A. diaperinus* (both Coleoptera) food products. Subclusters differentiated raw food products (flour) from processed food products (pasta, crackers and protein bars): flour made by the two insects of the Coleoptera order (i.e., *T. molitor* and *A. diaperinu*s) formed a distinct cluster that separated pasta and crackers samples based on the same insects. Moreover, same food products constituted by different insects can be distinguished by family abundances in the heatmap: *A. diaperinus* pasta clusterized separately from *T. molitor* pasta. Conversely, protein bars composed by the same insect (*A. domesticus*), but produced by different companies, are scattered in two different clusters, as also shown by microbial diversity analysis represented in the PCoA plot (Table S4).

### 3.4 Core microbiota

Core microbiota preliminary analyses revealed the taxa shared by at least 70% and the 100% of samples of the category representing the insect used in the food products. Venn diagram, calculated from core microbiota results of the most conserved taxa (100% of samples per insect), highlighted the presence of unique and shared taxa considering insect species used in the food products analysed (Fig 4).

**Fig 4.**
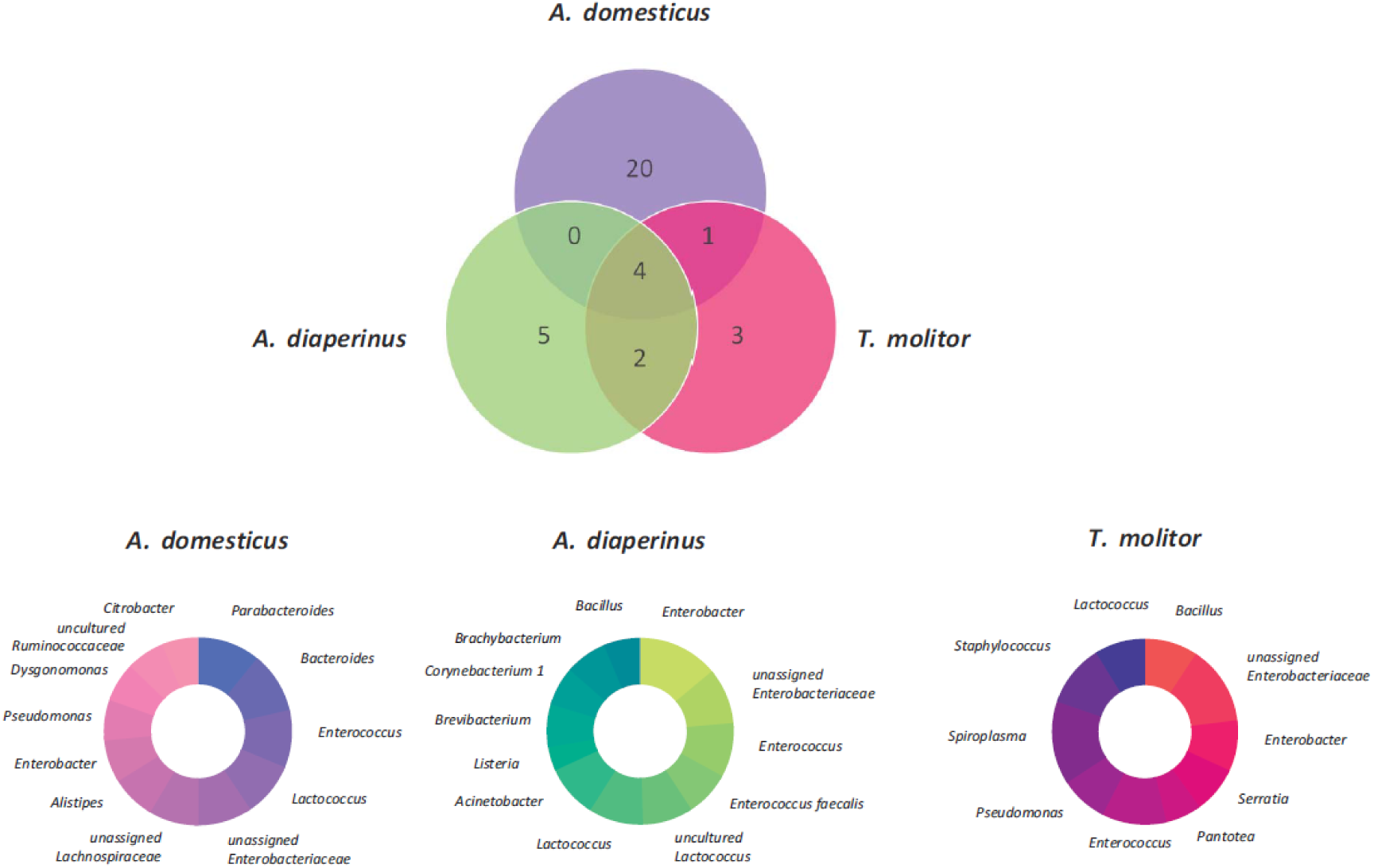
Venn diagram and donuts charts of *A. domesticus, A. diaperinus*, and *T. molitor* core microbial composition. Venn diagram in the upper part of the figure shows shared and unique taxa per insect. Taxa identified through core microbiota analysis are reported in the lower part of the figure. We considered the taxa found in 100% of the samples. In the case of *A. domesticus* and *A. diaperinus* the first twelve hits are reported, according to the frequency values listed in S4 Table.

In the case of *T. molitor*-based food products, we observed a core microbiota constituted by 21 taxa shared among > 70% of the samples and 10 taxa shared by all the samples. The 10 most conserved taxa (100% of samples) belonged to uncultured *Spiroplasma* sp., a taxon from Enterobacteriaceae family, *Enterococcus, Staphylococcus, Enterobacter*, uncultured *Lactococcus, Pseudomonas, Bacillus, Serratia* and *Pantotea* (S6 Table).

On the other hand, *A. diaperinus*-based food products showed 14 shared taxa, both subsampling the 70% of samples or considering all the samples, indicating a highly conserved core microbiota. In contrast to *T. molitor*-based products, we reported the presence not only of *Enterococcus, Staphylococcus, Enterobacter, Lactococcus*, but also of *Enterococcus faecalis, Listeria, Brevibacterium, Corynebacterium, Brachybacterium, Acinetobacter*, and *Bacillus pumilus.* We reported as well the absence of *Spiroplasma, Pseudomonas, Serratia* and *Pantotea*.

Considering *A. domesticus*-based food products, all the samples shared 29 taxa, and 44 taxa are shared by 70% of samples. Among these, all the samples reported the presence of bacteria belonging to the family Lachnospiraceae and to the genus *Parabateroides* (Family: Tannerellaceae).

Venn diagram analysis showed that, if four genera are shared among all the samples (a genus belonging to Enterobacteriaceae family, Lactococcus, Enterobacter, Enterococcus), 28 genera were unique considering the insect species. In particular, twenty genera were exclusively detected in all the samples of *A. domesticus*-based food products, and, among them, the three most abundant were Parabacteroides, Bacteroides, and a genus belonging to Lachnospiraceae family (see S6 Table for the complete list), thus confirming the explorative analyses described in the previous section. *Brevibacterium, Acinetobacter, Brachybacterium, Listeria*, and *Corynebacterium* were the genera unique for *A. diaperinus*-based food products, whereas *T. molitor*-based food products showed as unique genera *Spiroplasma, Pantoea*, and *Serratia*.

## 4. Discussion

In this study, we characterized through the application of HTS techniques the microbial composition of insect-based food products made of *A. domesticus, T. molitor*, and *A. diaperinus*, purchased via e-commerce. We selected both raw and processed food products, considering the availability on the market, from different selling companies.

Our preliminary data revealed that a small number of prevalent bacteria formed a “core microbiota” for each insect, which can potentially be used as biomarkers to identify insect ingredient origin in food products.

A recent study based on more than 20 samples per rearing condition analysed (plus 5 controls) (Cambon et al., 2018) showed that, although the relative abundances of some of the members of the microbiota are affected by rearing changes, a resident microbiota in *T. molitor* gut exists, thus supporting our hypothesis tested with core microbiota analysis. In particular, this study identified a resident *T. molitor* microbiota consisting of *Pseudomonas, Serratia* and genera belonging to the Enterobacteriaceae family. Noteworthy, this evidence is in accordance with the data we obtained in our study, as a further confirmation of our results. If there was a significant insect component, the core microbiota would reflect the physiology of the organisms, the diet and rearing conditions. By contrast, if the level of food processing affected the microbiota, the organism could be difficult to identify searching for a microbial signature. Nevertheless, we identified shared features constituting the core microbiota of specific insects. In addition to that, despite the processing level, we found exclusive taxa in all the samples of specific insects. Noteworthy, our results showed that in *A. domesticus* processed food (i.e. protein bars and crackers) microbiota is composed by a robust core of microorganisms that is conserved and is similar in composition to what was reported in other studies on raw food (i.e. fresh crickets): Vandeweyer and colleagues (Vandeweyer, Crauwels, Lievens, Van Campenhout, 2017) showed that *A. domesticus* is abundantly colonised by (Para)bacteroides species (Johnson, Moore, Moore, 2009), confirming the first two hits we obtained through core microbiota analysis.

Interestingly, in this study *A. domesticus* core microbiota harbured bacteria belonging to Lachnospiraceae family too. This evidence may prove beneficial when edible insects will be introduced in the western diet and it is worth further studies: Lachnospiraceae are found, among others, in our digestive tract and are involved in fiber digestion. The exposure to antibiotics (such as β-lactam antibiotics and fluoroquinolones) eliminates Lachnospiraceae from gut microbiota. This lead to the gut becoming a prime target for opportunistic infections such as the one caused by *Clostridium difficile*, but restoring Lachnospiraceae into the intestines of infected patients has been shown to help cure *C. difficile* infections (Lagier et al., 2012; Segata et al., 2012; Song et al., 2013; Seekatz, 2018). It is conceivable that in processed food we found only DNA and not viable cells and more investigations are needed, also focusing on prebiotic effects. In a recent study, the impact of an insect-based diet (cricket) on the human gut microbiota revealed increased levels of *Bifidobacterium animalis*. This could be due to cricket chitin which may function as a prebiotic (Stull et al., 2018). *T. molitor* flour in *in vitro* fecal models promoted the growth of Bacteroidaceae and Prevotellaceae, but not of *Clostridium histolyticum* group or Desulfovibrionales and Desulfuromonales (Carvallo et al., 2019).

On the other hand, exclusively all the samples based on *Tenebrio molitor* source are dominated by Spiroplasmataceae family (Phylum: Tenericutes; Class: Mollicutes), in particular bacteria belonging to *Spiroplasma* genus. *Spiroplasma* are a group of small bacteria without cell walls and share simple metabolism, parasitic lifestyle, and small genome (about 1 Mb).

*Spiroplasma* are found in the gut or hemolymph of insects where they can act as endosymbionts, impacting on host reproduction or host defence system. These findings are consistent with studies on fresh mealworm larvae (Vandeweyer, Crauwels, Lievens, Van Campenhout, 2017) deriving from different companies. They reported differences in the bacterial community composition that were higher in mealworms than in crickets, considering different rearing companies and production cycles.

To better disentangle these dynamics, Osimani and colleagues (Osimani et al., 2018) tested in laboratory conditions the microbiota changes of *Tenebrio molitor* rearing, “from feed to frass”: if wheatmeal showed low microbial contamination, both larvae and frass were characterized by *Enterobacter* spp., *Erwinia* spp., *Enterococcus* spp. and *Lactococcus* spp. as dominant genera. Entomoplasmatales (including *Spiroplasma* spp.) constituted a major fraction of the microbiota of larvae depending on the batch analysed and therefore suggesting that other unaccounted variables have a role in this.

*A. diaperinus* samples are dominated by *Enterobacter*, both flour and pasta, produced by different companies. These findings are in agreement with previous studies on fresh larvae (Wynants et al., 2017) and minced meat-like products (Stoops et al., 2016). *A. diaperinus*-based pasta clustered separately from flour samples made of the same insect, but in the same main cluster including food products belonging to Coleoptera. A similar behaviour can be seen in the case of *T. molitor* pasta and flour samples.

With regard to food safety, it is worth mentioning the presence, considering the 20 most abundant bacteria classified at the genus level, of sequences assigned to *Bacillus* in most of the samples (80%). The capacity to form endospore, resistant to heat and desiccation, deserve attention even if there is no confirmation of viability assay.

With the increasing availability of insect-based processed food products in the market, including a higher number of samples in the analyses will help in disentangling the microbial dynamics behind food processing, and allowing the food products traceability at a finer scale. Overall, our results showed that insect-based food products cluster based on their microbial signature. Even in the case of processed food in which there is more than one constituent (i.e., plant ingredients, see Table 1) that could interfere with its microbial contribution in the clustering process, we identified a shared pattern highlighted by core microbiota analysis and unique taxa that can be used as biomarkers. We also showed that differences exist in comparing raw vs processed food considering both qualitative and quantitative metrics. Recent studies (Bruno et al., 2019) reported the possibility to track the composition of plant processed food despite critical issues mostly deriving from the starting composition (i.e., variable complexity in taxa composition) of the sample itself and the different processing level (i.e., high or low DNA degradation). Other studies (Garofalo et al., 2017), investigating the microbial composition of commercial food products based on insects, never explored if any variability can be correlated with highly processed food such as pasta, crackers or protein bars. Our data clearly showed that processed food can be analysed searching for a microbial signature and that raw food products (i.e., flours) had a significant different microbiota compared to the processed ones (i.e., pasta, crackers and protein bars), even if maintaining unchanged a core of bacteria.

Highly processed food products represent one of the challenges of food traceability because of DNA degradation during food processing and, therefore, the limits in applying the common DNA barcoding techniques. Thus, DNA metabarcoding, based on HTS techniques combined with powerful tools for data analysis, can provide new perspectives for unveiling the composition of processed food, to retrace food origin and food quality control (Bruno et al., 2019; Parente et al., 2019; De Filippis, Parente & Ercolini, 2019).

The identification of a microbial signature for traceability purposes was suggested also by forensic scientists as natural consequence of the application of HTS technologies in a wider perspective (Bishop, 2019): with the globalisation of trade, food traceability is a hot topic and identifying a microbial signature in these products can provide a deeper insight into the “food ecosystem” (Galimberti et al., 2015; Bokulich, Lewis, Boundy-Mills, Mills, 2016; Galimberti et al., 2019; Parente et al., 2019).

## 5. Conclusions and future perspectives

The application of high-throughput molecular techniques coupled with bioinformatic analyses allowed us to detect and identify the diversity of microbial community in raw and processed novel food products available on e-commerce. We are now facing a striking imbalance between available technologies and knowledge gaps on “food ecosystem”: especially in the case of insect flour and insect based products, we should consider the whole food production chain, taking into consideration that the microbial communities inhabiting surfaces, interacting with foods and being part of food themselves are influenced all along the supply chain, from rearing, in the case of insects, to the final processed product. HTS approach is a valuable tool to protect food quality and safety as routine monitoring analysis, from the identification of insect microbiota along the food production processing chain and characterization of the raw ingredients to the final processed food products. This study shows the value of the application of HTS analysis for unveiling the composition of carried over insect microbiota in processed food containing insect ingredients. This tool can be applied in a wider range of food products to improve food source traceability and food quality control. Further studies are needed to improve our knowledge on the influence of rearing conditions and processing on the edible insect associated with the microbiota.

## Acknowledgements

The authors are grateful to Valerio Mezzasalma for assistance during experimental procedures and manuscript preparation and to Filippo Bargero for English revision. Thanks also to Simone Bosaglia and FlatIcon community for graphic contribution.

## Appendix A. Supplementary data

S1 Table. Insects used in the processed food analysed in this study.

S2 Table. List of primer pairs used for DNA barcoding and metabarcoding analyses.

S3 Table. Results of alpha microbial diversity.

S4 Table. Results of beta microbial diversity.

S5 Table. Microbial relative abundances (rank: Family)

S6 Table. Results of core microbiota analysis.

