## Supplementary material for "Tasting the differences: microbiota analysis of different insect-based novel food"

#### *Short title: Microbiota signature in insect-based food*

Jessica Frigerio[1¶] , Giulia Agostinetto[2¶], Andrea Galimberti[2], Fabrizio De Mattia[1], Massimo Labra[2], Antonia Bruno[2*]

[1]FEM2-Ambiente, Piazza della Scienza 2, I-20126 Milano, Italy;
[2]Zooplantlab, Department of Biotechnology and Biosciences, University of Milano-Bicocca, Piazza della Scienza 2, I-20126 Milano, Italy;

* Corresponding author


[¶]These authors contributed equally to this work

#### S1 Table. Insects used in the processed food analysed in this study.

| **Scientific name** | *Acheta domesticus* (Linnaeus, 1758) | *Tenebrio molitor* (Linnaeus, 1758) | *Alphitobius diaperinus* (Panzer, 1797) |
|---|---|---|---|
| **Taxonomy** | Order: Orthoptera Family Gryllidae | Order: Coleoptera Family: Tenebrionidae. | Order: Coleoptera Family: Tenebrionidae |
| **Description** | House cricket, native to Southwest Asia, widespread in tropical and temperate zones. Species is native to most of the European countries. | Known as mealworm. It has a cosmopolitan distribution, being common in Europe, as a pest of grain storages. | Known as lesser mealworm or litter beetle. It has a cosmopolitan distribution, being common in Europe, as a pest of grain storages and poultry farms. |
| **Growth** | Adults grow up to 20-22 mm, both sexes are fully winged. Adult females are slightly bigger with prominent ovipositor protruding from the abdomen. Crickets are greyish yellow in color. | The adult beetles are up to 15-18 mm long. It is shiny black or brown with reddish brown elytra.<br>The eggs are oval, whitish, about 1.5 mm long.<br>The larvae resemble larvae of other mealworms, at the final stage measuring up to 25 mm in length. | The adult beetles are 6 mm long, oval. It is shiny black or brown with reddish brown elytra. Color is variable among individuals and subpopulations and changing with age. The antennae are paler at the tips and are covered in tiny, yellowish hairs. The elytra have shallow longitudinal grooves. The eggs are narrow, whitish, about 1.5 mm long.<br>The larvae resemble larvae of other mealworms, at the final stage measuring up to 11 mm in length. |

| | | | |
|---|---|---|---|
| **Incubation period (days from egg-laying to hatch)** | 11 | 10-12 | 10-12 |
| **Time to maturity (days from hatch to max body weight)** | 32-49 | 280-400 | 280-400 |
| **Resistance** | Species is resistant to environmental conditions, and is very productive in mass culture, tolerating high population densities. The species is however very susceptible to the Cricket Paralysis Virus. | Species is resistant to environmental conditions, and is very productive in mass culture, tolerating high population densities. | Species is resistant to environmental conditions and is very productive in mass culture. |
| **Protein and fat content** | Protein content in larvae and imagines varies from 60 to 70% (d.m.), with a fat content of 20-25 % (d.m.) | Protein content in larvae varies from 50 to 65% (d.m.), with a fat content of 30-40 % (d.m.) highly depending on the feed and rearing conditions. | Protein content in larvae varies from 50 to 65% (d.m.), with fat content of 30-40 % (d.m.) highly depending on the feed and rearing conditions. |

Adapted from http://ipiff.org/wp-content/uploads/2019/03/IPIFF_Guide_A4_2019-v5-separate.pdf

#### S2 Table. List of primer pairs used for DNA barcoding and metabarcoding analyses.

| Primer name | 5'-3' | Barcode locus | Reference |
|---|---|---|---|
| LCO1490 | GGTCAACAAATCATAAAGATATTGG | COI | [17] |
| HC02198 | TAAACTTCAGGGTGACCAAAAAATCA | | |
| 340F | CTACGGGNGGCWGCAG | 16S | [19] |
| 806R | GACTACHVGGGTATCTAATCC | | |

#### S3 Table. Results of alpha microbial diversity.

**Pairwise comparison of ASVs counts between samples among to the same insect (*A. domesticus* n=12; *A. diaperinus* n=9; *T. molitor* n=15)**

| Group 1 | Group 2 | H | P-value | Q-value |
|---|---|---|---|---|

| | | | | |
|---|---|---|---|---|
| *A. domesticus* | *A. diaperinus* | 14.78 | < 0.01 | < 0.01 |
| *A. domesticus* | *T. molitor* | 5.83 | 0.015 | 0.015 |
| *A. diaperinus* | *T. molitor* | 14.82 | < 0.01 | < 0.01 |

**Pairwise comparison of Shannon index between samples among to the same insect (*A. domesticus* n=12; *A. diaperinus* n=9; *T. molitor* n=15)**

| Group 1 | Group 2 | H | P-value | Q-value |
|---|---|---|---|---|
| *A. domesticus* | *A. diaperinus* | 14.72 | < 0.01 | < 0.01 |
| *A. domesticus* | *T. molitor* | 19.28 | < 0.01 | < 0.01 |
| *A. diaperinus* | *T. molitor* | 16.20 | < 0.01 | < 0.01 |

#### S4 Table. Results of beta microbial diversity.

**PCoA Emperor plots based on the Bray-Curtis metric. Food samples were compared based on insect order (red: Coleoptera; blue: Orthoptera) and insect species (sphere: *A. domesticus*; triangle: *T. molitor*; square: *A. diaperinus*).**

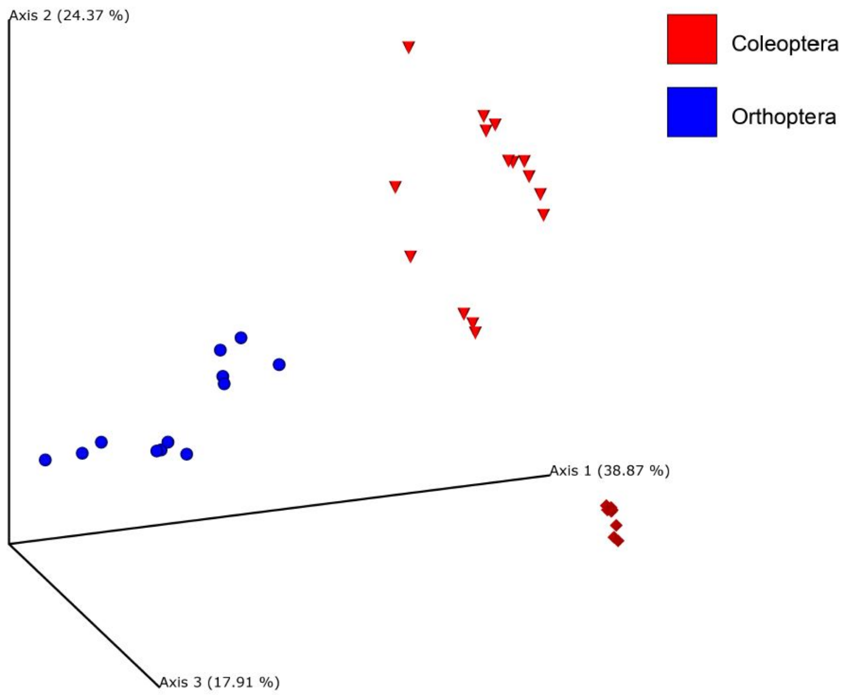

| **Raw materials (flour, n=12) vs food products (crackers, pasta, protein bars; n=24)** | **Pseudo-F** | **P-value** |
|---|---|---|
| Jaccard | 5.80 | 0.001 |
| Unweighted UniFrac | 5.40 | 0.001 |
| Bray-Curtis | 6.38 | 0.001 |
| Weighted UniFrac | 8.21 | 0.002 |

| **Differences among samples belong to different insects (*A. domesticus* n=12; *A. diaperinus* n=9; *T. molitor* n=15)** | **Pseudo-F** | **P-value** |
|---|---|---|
| Jaccard | 10.39 | 0.001 |
| Unweighted UniFrac | 11.37 | 0.001 |
| Bray-Curtis | 16.87 | 0.001 |
| Weighted UniFrac | 25.63 | 0.001 |

**Pairwise comparisons to test differences among samples belong to different insects (*A.domesticus* n=12; *A.diaperinus* n=9; *T.molitor* n=15).**

| **Insects** | **Group 1** | **Group 2** | **Pseudo-F** | **P-value** | **Q-value** |
|---|---|---|---|---|---|
| Jaccard | *A. domesticus* | *A. diaperinus* | 11.81 | 0.001 | 0.001 |
| | *A. domesticus* | *T. molitor* | 8.30 | 0.001 | 0.001 |
| | *A. diaperinus* | *T. molitor* | 12.35 | 0.001 | 0.001 |
| Unweighted UniFrac | *A. domesticus* | *A. diaperinus* | 14.51 | 0.001 | 0.001 |
| | *A. domesticus* | *T. molitor* | 12.27 | 0.001 | 0.001 |
| | *A. diaperinus* | *T. molitor* | 7.43 | 0.001 | 0.001 |
| Bray-Curtis | *A. domesticus* | *A. diaperinus* | 16.20 | 0.001 | 0.001 |
| | *A. domesticus* | *T. molitor* | 12.51 | 0.001 | 0.001 |
| | *A. diaperinus* | *T. molitor* | 25.95 | 0.001 | 0.001 |
| Weighted UniFrac | *A. domesticus* | *A. diaperinus* | 29.20 | 0.001 | 0.001 |
| | *A. domesticus* | *T. molitor* | 25.16 | 0.001 | 0.001 |

| | | | | | |
|---|---|---|---|---|---|
| | *A. diaperinus* | *T. molitor* | 21.19 | 0.001 | 0.001 |

### Supplementary S5

| | T.molitor_flour | T.molitor_flour | T.molitor_flour | T.molitor_flour | T.molitor_flour | T.molitor_flour | T.molitor_flour | T.molitor_flour | T.molitor_flour | A. diaperinus_flour | A. diaperinus_flour | A. diaperinus_flour | A. diaperinus_pasta | A. diaperinus_pasta | A. diaperinus_pasta | A. diaperinus_pasta | A. diaperinus_pasta | A. diaperinus_pasta | T.molitor_pasta | T.molitor_pasta | T.molitor_pasta | A. domesticus_cracker | A. domesticus_cracker | A. domesticus_cracker | T.molitor_cracker | T.molitor_cracker | T.molitor_cracker | A. domesticus_bar | A. domesticus_bar | A. domesticus_bar | A. domesticus_bar | A. domesticus_bar | A. domesticus_bar | A. domesticus_bar | A. domesticus_bar | A. domesticus_bar | | |
|---|---|---|---|---|---|---|---|---|---|---|---|---|---|---|---|---|---|---|---|---|---|---|---|---|---|---|---|---|---|---|---|---|---|---|---|---|---|---|
| taxa | R_001 | R_001 | R_001 | R_002 | R_002 | R_002 | R_003 | R_003 | R_003 | R_004 | R_004 | R_004 | FP_005 | FP_005 | FP_005 | FP_006 | FP_006 | FP_006 | FP_007 | FP_007 | FP_007 | FP_008 | FP_008 | FP_008 | FP_009 | FP_009 | FP_009 | FP_010 | FP_010 | FP_010 | FP_011 | FP_011 | FP_011 | FP_012 | FP_012 | FP_012 | SUM | % |
| D_0__Bacteria;D_1__Proteobacteria;D_2__Gammaproteobacteria;D_3__Enterobacteriales;D_4__Enterobacteriaceae | 15949 | 30355 | 24500 | 13013 | 15147 | 19227 | 22166 | 17985 | 35623 | 133873 | 116902 | 48649 | 4664 | 3468 | 4501 | 8194 | 7575 | 5239 | 239 | 744 | 409 | 289 | 197 | 343 | 300 | 463 | 1721 | 235 | 256 | 211 | 594 | 784 | 182 | 1304 | 861 | 1731 | 537893 | 45.1837 |
| D_0__Bacteria;D_1__Tenericutes;D_2__Mollicutes;D_3__Entomoplasmatales;D_4__Spiroplasmataceae | 32256 | 45829 | 36616 | 12080 | 15598 | 43262 | 36834 | 38357 | 41932 | 0 | 0 | 4 | 0 | 0 | 0 | 10 | 6 | 4 | 30 | 87 | 65 | 26 | 0 | 8 | 1024 | 418 | 509 | 0 | 0 | 0 | 0 | 0 | 0 | 0 | 0 | 0 | 304955 | 25.61661 |
| D_0__Bacteria;D_1__Firmicutes;D_2__Bacilli;D_3__Bacillales;D_4__Staphylococcaceae | 5106 | 4363 | 3355 | 1485 | 1248 | 2332 | 2175 | 1815 | 3768 | 2214 | 1149 | 640 | 216 | 167 | 298 | 579 | 519 | 410 | 20 | 20 | 21 | 352 | 189 | 4 | 94 | 197 | 1023 | 0 | 0 | 0 | 0 | 0 | 0 | 5301 | 6689 | 4336 | 50085 | 4.207204 |
| D_0__Bacteria;D_1__Firmicutes;D_2__Bacilli;D_3__Lactobacillales;D_4__Streptococcaceae | 4160 | 3719 | 3689 | 2233 | 2115 | 2518 | 1916 | 1857 | 3541 | 8074 | 3175 | 1837 | 303 | 250 | 325 | 560 | 597 | 415 | 96 | 282 | 141 | 230 | 102 | 78 | 29 | 15 | 1035 | 11 | 12 | 18 | 29 | 33 | 8 | 1628 | 1978 | 1285 | 48294 | 4.056758 |
| D_0__Bacteria;D_1__Firmicutes;D_2__Bacilli;D_3__Lactobacillales;D_4__Enterococcaceae | 4635 | 3649 | 4095 | 1466 | 1214 | 2660 | 2515 | 2461 | 4814 | 4975 | 2562 | 1265 | 459 | 278 | 361 | 604 | 563 | 390 | 184 | 351 | 222 | 45 | 75 | 167 | 30 | 0 | 0 | 50 | 67 | 72 | 115 | 147 | 31 | 1964 | 2474 | 1526 | 46486 | 3.904884 |
| D_0__Bacteria;D_1__Firmicutes;D_2__Bacilli;D_3__Bacillales;D_4__Planococcaceae | 0 | 0 | 0 | 0 | 0 | 0 | 0 | 0 | 0 | 14 | 0 | 8 | 0 | 0 | 0 | 0 | 0 | 0 | 0 | 0 | 0 | 0 | 0 | 0 | 91 | 131 | 0 | 0 | 0 | 0 | 0 | 0 | 0 | 11038 | 11993 | 9217 | 32492 | 2.72937 |
| D_0__Bacteria;D_1__Firmicutes;D_2__Bacilli;D_3__Lactobacillales;D_4__Lactobacillaceae | 4586 | 4000 | 4535 | 2130 | 2026 | 2693 | 2581 | 2208 | 4761 | 60 | 26 | 13 | 0 | 0 | 0 | 5 | 0 | 0 | 0 | 5 | 2 | 22 | 53 | 16 | 17 | 5 | 260 | 0 | 0 | 0 | 4 | 5 | 0 | 47 | 11 | 2 | 30073 | 2.526171 |
| D_0__Bacteria;D_1__Firmicutes;D_2__Bacilli;D_3__Bacillales;D_4__Bacillaceae | 207 | 215 | 207 | 96 | 26 | 117 | 93 | 109 | 126 | 170 | 73 | 12 | 9 | 0 | 13 | 15 | 6 | 0 | 169 | 459 | 317 | 1792 | 275 | 32 | 315 | 695 | 2212 | 0 | 0 | 0 | 20 | 20 | 16 | 4708 | 7361 | 5043 | 24928 | 2.093984 |
| D_0__Bacteria;D_1__Bacteroidetes;D_2__Bacteroidia;D_3__Bacteroidales;D_4__Tannerellaceae | 0 | 0 | 0 | 0 | 0 | 0 | 0 | 0 | 0 | 0 | 0 | 0 | 0 | 0 | 0 | 0 | 0 | 0 | 0 | 0 | 0 | 402 | 143 | 2114 | 0 | 0 | 0 | 244 | 531 | 340 | 1139 | 1122 | 262 | 5333 | 2843 | 4050 | 18523 | 1.555956 |
| D_0__Bacteria;D_1__Firmicutes;D_2__Bacilli;D_3__Lactobacillales;D_4__Leuconostocaceae | 1885 | 1852 | 1571 | 693 | 804 | 1080 | 960 | 945 | 2078 | 41 | 25 | 9 | 0 | 0 | 0 | 6 | 5 | 0 | 0 | 7 | 0 | 28 | 0 | 15 | 7 | 0 | 0 | 0 | 0 | 0 | 6 | 0 | 0 | 167 | 295 | 148 | 12627 | 1.060684 |
| D_0__Bacteria;D_1__Firmicutes;D_2__Bacilli;D_3__Bacillales;D_4__Family XII | 0 | 0 | 0 | 0 | 0 | 0 | 0 | 0 | 0 | 0 | 0 | 0 | 0 | 0 | 0 | 0 | 0 | 0 | 0 | 0 | 0 | 16 | 0 | 0 | 0 | 0 | 0 | 0 | 0 | 0 | 0 | 0 | 0 | 3402 | 5229 | 3340 | 11987 | 1.006923 |
| D_0__Bacteria;D_1__Bacteroidetes;D_2__Bacteroidia;D_3__Bacteroidales;D_4__Bacteroidaceae | 0 | 0 | 0 | 0 | 0 | 0 | 0 | 0 | 0 | 0 | 7 | 0 | 0 | 0 | 0 | 0 | 0 | 0 | 0 | 0 | 0 | 286 | 63 | 648 | 0 | 0 | 0 | 40 | 130 | 78 | 416 | 300 | 47 | 3174 | 1768 | 2304 | 9261 | 0.777936 |
| D_0__Bacteria;D_1__Proteobacteria;D_2__Gammaproteobacteria;D_3__Pseudomonadales;D_4__Pseudomonadaceae | 785 | 1239 | 901 | 573 | 295 | 905 | 370 | 238 | 632 | 42 | 0 | 12 | 6 | 0 | 0 | 16 | 10 | 8 | 37 | 116 | 33 | 44 | 49 | 92 | 155 | 118 | 1678 | 7 | 117 | 26 | 51 | 60 | 16 | 175 | 232 | 137 | 9175 | 0.770712 |
| D_0__Bacteria;D_1__Firmicutes;D_2__Clostridia;D_3__Clostridiales;D_4__Clostridiaceae 1 | 600 | 800 | 690 | 1121 | 1352 | 962 | 515 | 530 | 591 | 0 | 0 | 0 | 0 | 0 | 0 | 0 | 0 | 0 | 0 | 0 | 0 | 0 | 0 | 0 | 0 | 0 | 0 | 0 | 0 | 0 | 0 | 0 | 0 | 0 | 0 | 0 | 7161 | 0.601533 |
| D_0__Bacteria;D_1__Actinobacteria;D_2__Actinobacteria;D_3__Micrococcales;D_4__Brevibacteriaceae | 860 | 1122 | 653 | 290 | 390 | 591 | 361 | 370 | 618 | 485 | 203 | 142 | 12 | 9 | 18 | 16 | 15 | 13 | 0 | 0 | 0 | 0 | 0 | 0 | 0 | 0 | 0 | 0 | 0 | 0 | 0 | 0 | 0 | 0 | 0 | 0 | 6168 | 0.51812 |
| D_0__Bacteria;D_1__Firmicutes;D_2__Bacilli;D_3__Bacillales;D_4__Listeriaceae | 490 | 291 | 620 | 290 | 139 | 299 | 345 | 263 | 452 | 954 | 300 | 149 | 21 | 9 | 15 | 16 | 37 | 10 | 0 | 0 | 0 | 0 | 0 | 0 | 12 | 0 | 416 | 0 | 0 | 0 | 0 | 0 | 0 | 0 | 0 | 0 | 5128 | 0.430759 |
| D_0__Bacteria;D_1__Proteobacteria;D_2__Gammaproteobacteria;D_3__Pseudomonadales;D_4__Moraxellaceae | 18 | 122 | 12 | 19 | 48 | 45 | 16 | 0 | 32 | 49 | 26 | 18 | 0 | 0 | 10 | 0 | 0 | 6 | 0 | 0 | 0 | 63 | 0 | 12 | 26 | 34 | 362 | 0 | 0 | 0 | 0 | 0 | 0 | 1173 | 1682 | 1224 | 4997 | 0.419754 |
| D_0__Bacteria;D_1__Actinobacteria;D_2__Actinobacteria;D_3__Corynebacteriales;D_4__Corynebacteriaceae | 244 | 700 | 140 | 90 | 120 | 143 | 55 | 39 | 115 | 736 | 397 | 253 | 10 | 8 | 12 | 15 | 14 | 11 | 0 | 0 | 0 | 141 | 120 | 28 | 44 | 31 | 429 | 0 | 0 | 0 | 0 | 0 | 0 | 9 | 0 | 0 | 3904 | 0.327941 |
| D_0__Bacteria;D_1__Firmicutes;D_2__Clostridia;D_3__Clostridiales;D_4__Lachnospiraceae | 5 | 14 | 0 | 3 | 0 | 0 | 237 | 330 | 0 | 0 | 0 | 0 | 0 | 0 | 0 | 0 | 0 | 0 | 0 | 3 | 0 | 228 | 122 | 352 | 0 | 34 | 92 | 116 | 134 | 80 | 628 | 397 | 84 | 450 | 195 | 179 | 3683 | 0.309377 |
| D_0__Bacteria;D_1__Actinobacteria;D_2__Actinobacteria;D_3__Micrococcales;D_4__Dermabacteraceae | 282 | 426 | 238 | 127 | 174 | 272 | 188 | 177 | 345 | 343 | 125 | 91 | 25 | 0 | 15 | 16 | 0 | 0 | 0 | 0 | 0 | 16 | 0 | 0 | 0 | 0 | 0 | 0 | 0 | 0 | 0 | 0 | 0 | 0 | 0 | 0 | 2860 | 0.240244 |
| D_0__Bacteria;D_1__Firmicutes;D_2__Clostridia;D_3__Clostridiales;D_4__Ruminococcaceae | 0 | 0 | 0 | 0 | 0 | 0 | 0 | 0 | 0 | 0 | 0 | 2 | 0 | 0 | 0 | 0 | 0 | 0 | 0 | 0 | 0 | 96 | 136 | 238 | 0 | 0 | 0 | 116 | 214 | 146 | 426 | 299 | 90 | 296 | 59 | 143 | 2261 | 0.189927 |
| D_0__Bacteria;D_1__Cyanobacteria;D_2__Oxyphotobacteria;D_3__Nostocales;D_4__Phormidiaceae | 0 | 0 | 0 | 0 | 0 | 0 | 0 | 0 | 0 | 0 | 0 | 8 | 0 | 0 | 0 | 0 | 0 | 0 | 0 | 0 | 0 | 0 | 0 | 0 | 13 | 0 | 2110 | 0 | 0 | 0 | 0 | 0 | 0 | 0 | 0 | 0 | 2131 | 0.179007 |
| D_0__Bacteria;D_1__Bacteroidetes;D_2__Bacteroidia;D_3__Bacteroidales;D_4__Rikenellaceae | 0 | 2 | 0 | 0 | 0 | 0 | 0 | 0 | 0 | 0 | 0 | 0 | 0 | 0 | 0 | 0 | 0 | 0 | 0 | 0 | 0 | 15 | 34 | 651 | 0 | 0 | 0 | 41 | 130 | 29 | 142 | 141 | 9 | 436 | 175 | 269 | 2074 | 0.174219 |
| D_0__Bacteria;D_1__Actinobacteria;D_2__Actinobacteria;D_3__Propionibacteriales;D_4__Propionibacteriaceae | 0 | 0 | 0 | 0 | 0 | 0 | 0 | 0 | 0 | 0 | 0 | 0 | 0 | 0 | 0 | 0 | 0 | 0 | 0 | 0 | 0 | 304 | 126 | 47 | 64 | 179 | 1147 | 0 | 0 | 0 | 0 | 0 | 0 | 6 | 6 | 7 | 1886 | 0.158426 |
| Unassigned;__;__;__;__ | 203 | 218 | 264 | 61 | 32 | 41 | 0 | 58 | 97 | 131 | 54 | 17 | 0 | 0 | 0 | 0 | 0 | 0 | 0 | 0 | 0 | 62 | 81 | 0 | 91 | 120 | 0 | 0 | 0 | 0 | 0 | 0 | 0 | 8 | 0 | 0 | 1538 | 0.129194 |
| D_0__Bacteria;D_1__Bacteroidetes;D_2__Bacteroidia;D_3__Bacteroidales;D_4__Dysgonomonadaceae | 8 | 0 | 9 | 0 | 0 | 0 | 0 | 6 | 0 | 0 | 4 | 0 | 0 | 0 | 0 | 0 | 0 | 0 | 0 | 0 | 0 | 0 | 6 | 403 | 0 | 0 | 0 | 29 | 30 | 23 | 149 | 151 | 10 | 251 | 68 | 75 | 1222 | 0.10265 |
| D_0__Bacteria;D_1__Firmicutes;D_2__Clostridia;D_3__Clostridiales;D_4__Peptostreptococcaceae | 139 | 130 | 170 | 55 | 44 | 88 | 87 | 80 | 203 | 0 | 0 | 0 | 0 | 0 | 0 | 0 | 0 | 0 | 0 | 0 | 0 | 0 | 0 | 0 | 0 | 0 | 0 | 0 | 0 | 0 | 0 | 0 | 0 | 0 | 0 | 0 | 996 | 0.083665 |
| D_0__Bacteria;D_1__Actinobacteria;D_2__Actinobacteria;D_3__Micrococcales;D_4__Micrococcaceae | 100 | 125 | 81 | 31 | 40 | 51 | 45 | 58 | 108 | 29 | 12 | 0 | 0 | 0 | 0 | 0 | 0 | 0 | 0 | 0 | 0 | 9 | 0 | 0 | 0 | 20 | 140 | 0 | 0 | 0 | 0 | 0 | 0 | 0 | 0 | 0 | 849 | 0.071317 |
| D_0__Bacteria;D_1__Proteobacteria;D_2__Deltaproteobacteria;D_3__Desulfovibrionales;D_4__Desulfovibrionaceae | 0 | 0 | 0 | 0 | 0 | 0 | 0 | 0 | 0 | 0 | 0 | 0 | 0 | 0 | 0 | 0 | 0 | 0 | 0 | 0 | 0 | 0 | 0 | 70 | 0 | 0 | 0 | 0 | 28 | 7 | 49 | 54 | 0 | 256 | 89 | 78 | 631 | 0.053005 |
| D_0__Bacteria;D_1__Proteobacteria;D_2__Gammaproteobacteria;D_3__Betaproteobacteriales;D_4__Neisseriaceae | 0 | 0 | 0 | 0 | 0 | 0 | 0 | 0 | 0 | 0 | 0 | 0 | 0 | 0 | 0 | 0 | 0 | 0 | 0 | 0 | 0 | 90 | 102 | 109 | 0 | 28 | 90 | 3 | 0 | 0 | 7 | 17 | 4 | 21 | 7 | 6 | 484 | 0.040657 |
| D_0__Bacteria;D_1__Bacteroidetes;D_2__Bacteroidia;D_3__Flavobacteriales;D_4__Weeksellaceae | 0 | 11 | 0 | 0 | 2 | 8 | 2 | 0 | 0 | 0 | 0 | 0 | 0 | 0 | 0 | 0 | 0 | 0 | 11 | 44 | 20 | 0 | 0 | 0 | 43 | 16 | 0 | 0 | 10 | 17 | 30 | 10 | 11 | 60 | 59 | 52 | 406 | 0.034105 |
| D_0__Bacteria;D_1__Firmicutes;D_2__Clostridia;D_3__Clostridiales;D_4__Clostridiales vadinBB60 group | 0 | 0 | 0 | 0 | 0 | 0 | 0 | 0 | 0 | 0 | 0 | 0 | 0 | 0 | 0 | 0 | 0 | 0 | 0 | 0 | 0 | 0 | 0 | 8 | 0 | 0 | 0 | 0 | 0 | 0 | 0 | 3 | 0 | 160 | 109 | 108 | 388 | 0.032592 |
| D_0__Bacteria;D_1__Proteobacteria;D_2__Gammaproteobacteria;D_3__Xanthomonadales;D_4__Xanthomonadaceae | 10 | 15 | 10 | 16 | 21 | 22 | 27 | 7 | 79 | 0 | 0 | 0 | 0 | 0 | 0 | 0 | 0 | 0 | 5 | 13 | 8 | 0 | 0 | 0 | 0 | 0 | 0 | 8 | 50 | 17 | 25 | 38 | 0 | 0 | 0 | 0 | 371 | 0.031164 |
| D_0__Bacteria;D_1__Firmicutes;D_2__Negativicutes;D_3__Selenomonadales;D_4__Acidaminococcaceae | 0 | 0 | 0 | 0 | 0 | 0 | 0 | 0 | 0 | 0 | 0 | 0 | 0 | 0 | 0 | 0 | 0 | 0 | 0 | 0 | 0 | 0 | 0 | 39 | 0 | 0 | 0 | 6 | 3 | 0 | 19 | 28 | 0 | 76 | 44 | 117 | 332 | 0.027888 |
| D_0__Bacteria;D_1__Firmicutes;D_2__Clostridia;D_3__Clostridiales;D_4__Peptococcaceae | 0 | 0 | 0 | 0 | 0 | 0 | 0 | 0 | 0 | 0 | 0 | 0 | 0 | 0 | 0 | 0 | 0 | 0 | 0 | 0 | 0 | 23 | 0 | 35 | 0 | 0 | 0 | 0 | 0 | 0 | 0 | 0 | 0 | 134 | 57 | 72 | 321 | 0.026964 |
| D_0__Bacteria;D_1__Verrucomicrobia;D_2__Verrucomicrobiae;D_3__Verrucomicrobiales;D_4__Akkermansiaceae | 0 | 0 | 0 | 0 | 0 | 0 | 0 | 0 | 0 | 0 | 0 | 0 | 0 | 0 | 0 | 0 | 0 | 0 | 0 | 0 | 0 | 13 | 4 | 31 | 0 | 0 | 0 | 3 | 0 | 11 | 15 | 14 | 0 | 103 | 29 | 58 | 281 | 0.023604 |
| D_0__Bacteria;D_1__Actinobacteria;D_2__Actinobacteria;D_3__Bifidobacteriales;D_4__Bifidobacteriaceae | 0 | 0 | 0 | 0 | 0 | 0 | 0 | 0 | 0 | 0 | 0 | 0 | 0 | 0 | 0 | 0 | 0 | 0 | 0 | 0 | 0 | 0 | 42 | 70 | 0 | 0 | 0 | 0 | 0 | 0 | 0 | 0 | 0 | 0 | 68 | 85 | 265 | 0.02226 |
| D_0__Bacteria;D_1__Firmicutes;D_2__Bacilli;D_3__Lactobacillales;D_4__Carnobacteriaceae | 0 | 37 | 0 | 0 | 0 | 0 | 0 | 0 | 0 | 0 | 0 | 0 | 0 | 0 | 0 | 0 | 0 | 0 | 0 | 0 | 0 | 18 | 0 | 29 | 0 | 18 | 0 | 0 | 0 | 0 | 0 | 0 | 0 | 49 | 65 | 37 | 253 | 0.021252 |
| D_0__Bacteria;D_1__Firmicutes;D_2__Bacilli;D_3__Bacillales;D_4__Thermoactinomycetaceae | 0 | 0 | 0 | 0 | 0 | 0 | 0 | 0 | 0 | 0 | 0 | 0 | 0 | 0 | 0 | 0 | 0 | 0 | 0 | 2 | 0 | 40 | 188 | 0 | 0 | 0 | 0 | 0 | 0 | 0 | 0 | 0 | 0 | 0 | 0 | 0 | 230 | 0.01932 |
| D_0__Bacteria;D_1__Proteobacteria;D_2__Gammaproteobacteria;D_3__Betaproteobacteriales;D_4__Burkholderiaceae | 12 | 0 | 0 | 0 | 0 | 0 | 0 | 0 | 0 | 0 | 0 | 0 | 0 | 0 | 0 | 0 | 0 | 0 | 0 | 0 | 0 | 23 | 0 | 0 | 0 | 0 | 0 | 0 | 0 | 0 | 0 | 0 | 0 | 32 | 80 | 79 | 226 | 0.018984 |
| D_0__Bacteria;D_1__Firmicutes;D_2__Erysipelotrichia;D_3__Erysipelotrichales;D_4__Erysipelotrichaceae | 0 | 0 | 0 | 0 | 0 | 0 | 0 | 0 | 0 | 0 | 0 | 2 | 0 | 0 | 0 | 3 | 0 | 0 | 0 | 0 | 0 | 16 | 8 | 9 | 0 | 0 | 0 | 0 | 7 | 11 | 9 | 28 | 10 | 68 | 32 | 21 | 224 | 0.018816 |
| D_0__Bacteria;D_1__Proteobacteria;D_2__Gammaproteobacteria;D_3__Aeromonadales;D_4__Aeromonadaceae | 6 | 67 | 126 | 13 | 0 | 0 | 0 | 0 | 0 | 0 | 0 | 0 | 0 | 0 | 0 | 0 | 0 | 0 | 0 | 0 | 0 | 0 | 0 | 0 | 0 | 0 | 0 | 0 | 0 | 0 | 0 | 0 | 0 | 0 | 0 | 0 | 212 | 0.017808 |
| D_0__Bacteria;D_1__Proteobacteria;D_2__Alphaproteobacteria;D_3__Rhizobiales;D_4__Rhizobiaceae | 0 | 0 | 0 | 0 | 0 | 0 | 0 | 0 | 0 | 0 | 0 | 0 | 0 | 0 | 0 | 0 | 0 | 0 | 33 | 55 | 43 | 0 | 0 | 0 | 8 | 72 | 0 | 0 | 0 | 0 | 0 | 0 | 0 | 0 | 0 | 0 | 211 | 0.017724 |
| D_0__Bacteria;D_1__Firmicutes;D_2__Clostridia;D_3__Clostridiales;D_4__Family XIII | 0 | 0 | 0 | 0 | 0 | 0 | 0 | 0 | 0 | 0 | 0 | 0 | 0 | 0 | 0 | 0 | 0 | 0 | 0 | 0 | 0 | 0 | 0 | 15 | 0 | 0 | 0 | 13 | 9 | 0 | 25 | 71 | 0 | 38 | 20 | 0 | 191 | 0.016044 |
| D_0__Bacteria;D_1__Proteobacteria;D_2__Alphaproteobacteria;D_3__Rhizobiales;D_4__Beijerinckiaceae | 0 | 0 | 0 | 0 | 0 | 0 | 0 | 0 | 0 | 0 | 0 | 0 | 0 | 0 | 0 | 0 | 0 | 0 | 31 | 78 | 51 | 0 | 0 | 0 | 0 | 0 | 20 | 0 | 0 | 0 | 0 | 0 | 0 | 0 | 0 | 0 | 180 | 0.01512 |
| D_0__Bacteria;D_1__Proteobacteria;D_2__Alphaproteobacteria;D_3__Sphingomonadales;D_4__Sphingomonadaceae | 0 | 0 | 0 | 0 | 0 | 0 | 0 | 0 | 0 | 0 | 0 | 0 | 0 | 0 | 0 | 0 | 6 | 0 | 22 | 54 | 62 | 0 | 0 | 0 | 9 | 0 | 0 | 0 | 0 | 0 | 0 | 0 | 0 | 0 | 13 | 9 | 175 | 0.0147 |
| D_0__Bacteria;D_1__Bacteroidetes;D_2__Bacteroidia;D_3__Bacteroidales;D_4__Paludibacteraceae | 0 | 0 | 0 | 0 | 0 | 0 | 0 | 0 | 0 | 0 | 0 | 0 | 0 | 0 | 0 | 0 | 0 | 0 | 0 | 0 | 0 | 0 | 0 | 25 | 0 | 0 | 0 | 6 | 32 | 0 | 22 | 12 | 0 | 46 | 14 | 11 | 168 | 0.014112 |
| D_0__Bacteria;D_1__Proteobacteria;D_2__Gammaproteobacteria;D_3__Oceanospirillales;D_4__Halomonadaceae | 0 | 0 | 19 | 0 | 18 | 41 | 0 | 0 | 9 | 0 | 0 | 0 | 0 | 0 | 0 | 0 | 0 | 0 | 0 | 0 | 0 | 0 | 0 | 0 | 0 | 0 | 0 | 0 | 0 | 0 | 31 | 0 | 13 | 26 | 3 | 0 | 160 | 0.01344 |
| D_0__Bacteria;D_1__Firmicutes;D_2__Bacilli;D_3__Bacillales;D_4__Paenibacillaceae | 0 | 0 | 0 | 0 | 0 | 0 | 0 | 0 | 0 | 9 | 0 | 0 | 0 | 0 | 0 | 2 | 0 | 0 | 0 | 0 | 0 | 0 | 0 | 0 | 0 | 0 | 0 | 0 | 0 | 0 | 0 | 0 | 0 | 42 | 85 | 0 | 138 | 0.011592 |
| D_0__Bacteria;D_1__Fusobacteria;D_2__Fusobacteriia;D_3__Fusobacteriales;D_4__Leptotrichiaceae | 0 | 0 | 0 | 0 | 0 | 0 | 0 | 0 | 0 | 0 | 0 | 0 | 0 | 0 | 0 | 0 | 0 | 0 | 8 | 68 | 41 | 0 | 0 | 0 | 12 | 0 | 0 | 0 | 0 | 0 | 0 | 0 | 0 | 0 | 0 | 0 | 129 | 0.010836 |
| D_0__Bacteria;D_1__Actinobacteria;D_2__Actinobacteria;D_3__Micrococcales;D_4__Microbacteriaceae | 11 | 0 | 0 | 0 | 10 | 8 | 0 | 0 | 0 | 0 | 0 | 0 | 0 | 0 | 0 | 0 | 0 | 0 | 13 | 37 | 30 | 0 | 0 | 0 | 0 | 0 | 0 | 0 | 0 | 0 | 0 | 0 | 0 | 0 | 0 | 0 | 109 | 0.009156 |
| D_0__Bacteria;D_1__Proteobacteria;D_2__Gammaproteobacteria;D_3__Alteromonadales;D_4__Shewanellaceae | 0 | 102 | 0 | 0 | 0 | 0 | 0 | 0 | 0 | 0 | 0 | 0 | 0 | 0 | 0 | 0 | 0 | 0 | 0 | 0 | 0 | 0 | 0 | 0 | 0 | 0 | 0 | 0 | 0 | 0 | 0 | 0 | 0 | 0 | 0 | 0 | 102 | 0.008568 |
| D_0__Bacteria;D_1__Fusobacteria;D_2__Fusobacteriia;D_3__Fusobacteriales;D_4__Fusobacteriaceae | 0 | 0 | 0 | 0 | 0 | 0 | 0 | 0 | 0 | 0 | 0 | 0 | 0 | 0 | 0 | 0 | 0 | 0 | 0 | 0 | 0 | 0 | 0 | 17 | 0 | 0 | 0 | 0 | 0 | 0 | 4 | 0 | 0 | 36 | 0 | 27 | 84 | 0.007056 |
| D_0__Bacteria;D_1__Proteobacteria;D_2__Alphaproteobacteria;D_3__Rhodospirillales;D_4__Magnetospirillaceae | 0 | 0 | 0 | 0 | 0 | 0 | 0 | 0 | 0 | 0 | 0 | 0 | 0 | 0 | 0 | 0 | 0 | 0 | 0 | 0 | 0 | 0 | 0 | 4 | 0 | 0 | 0 | 4 | 15 | 4 | 11 | 10 | 0 | 10 | 3 | 11 | 72 | 0.006048 |
| D_0__Bacteria;D_1__Proteobacteria;D_2__Alphaproteobacteria;D_3__Acetobacterales;D_4__Acetobacteraceae | 15 | 6 | 5 | 0 | 0 | 0 | 0 | 0 | 6 | 0 | 0 | 0 | 0 | 0 | 0 | 0 | 0 | 0 | 0 | 0 | 0 | 21 | 11 | 0 | 0 | 0 | 0 | 0 | 0 | 0 | 0 | 0 | 0 | 0 | 0 | 0 | 64 | 0.005376 |
| D_0__Bacteria;D_1__Firmicutes;D_2__Clostridia;D_3__Clostridiales;D_4__Christensenellaceae | 0 | 0 | 0 | 0 | 0 | 0 | 0 | 0 | 0 | 0 | 0 | 0 | 0 | 0 | 0 | 0 | 0 | 0 | 0 | 0 | 0 | 15 | 0 | 7 | 0 | 0 | 0 | 0 | 4 | 0 | 13 | 13 | 2 | 0 | 6 | 0 | 60 | 0.00504 |
| D_0__Bacteria;D_1__Proteobacteria;D_2__Gammaproteobacteria;D_3__Pasteurellales;D_4__Pasteurellaceae | 0 | 0 | 0 | 0 | 0 | 0 | 0 | 0 | 0 | 0 | 0 | 0 | 0 | 0 | 0 | 0 | 0 | 0 | 0 | 0 | 0 | 0 | 0 | 0 | 34 | 23 | 0 | 0 | 0 | 0 | 0 | 0 | 0 | 0 | 0 | 0 | 57 | 0.004788 |
| D_0__Bacteria;D_1__Actinobacteria;D_2__Actinobacteria;D_3__Pseudonocardiales;D_4__Pseudonocardiaceae | 0 | 0 | 0 | 0 | 0 | 0 | 0 | 0 | 0 | 0 | 0 | 0 | 0 | 0 | 0 | 0 | 0 | 0 | 0 | 0 | 0 | 18 | 0 | 5 | 0 | 0 | 0 | 0 | 0 | 0 | 0 | 0 | 0 | 5 | 12 | 8 | 48 | 0.004032 |
| D_0__Bacteria;D_1__Actinobacteria;D_2__Actinobacteria;D_3__Corynebacteriales;D_4__Dietziaceae | 0 | 0 | 0 | 0 | 0 | 0 | 0 | 0 | 0 | 25 | 18 | 0 | 0 | 0 | 0 | 0 | 0 | 0 | 0 | 0 | 0 | 0 | 0 | 0 | 0 | 0 | 0 | 0 | 0 | 0 | 0 | 0 | 0 | 0 | 0 | 0 | 43 | 0.003612 |
| D_0__Bacteria;D_1__Bacteroidetes;D_2__Bacteroidia;D_3__Bacteroidales;D_4__Porphyromonadaceae | 0 | 0 | 0 | 0 | 0 | 0 | 0 | 0 | 0 | 0 | 0 | 0 | 0 | 0 | 0 | 0 | 0 | 0 | 0 | 0 | 0 | 0 | 0 | 0 | 0 | 0 | 0 | 3 | 0 | 0 | 26 | 14 | 0 | 0 | 0 | 0 | 43 | 0.003612 |
| D_0__Bacteria;D_1__Bacteroidetes;D_2__Bacteroidia;D_3__Sphingobacteriales;D_4__Sphingobacteriaceae | 0 | 0 | 0 | 0 | 20 | 16 | 0 | 0 | 7 | 0 | 0 | 0 | 0 | 0 | 0 | 0 | 0 | 0 | 0 | 0 | 0 | 0 | 0 | 0 | 0 | 0 | 0 | 0 | 0 | 0 | 0 | 0 | 0 | 0 | 0 | 0 | 43 | 0.003612 |
| D_0__Bacteria;D_1__Tenericutes;D_2__Mollicutes;D_3__Mycoplasmatales;D_4__Mycoplasmataceae | 0 | 0 | 0 | 0 | 0 | 0 | 0 | 0 | 0 | 14 | 7 | 0 | 0 | 0 | 0 | 5 | 7 | 5 | 0 | 0 | 0 | 0 | 0 | 0 | 0 | 0 | 0 | 0 | 0 | 0 | 0 | 0 | 0 | 0 | 0 | 0 | 38 | 0.003192 |
| D_0__Bacteria;D_1__Proteobacteria;D_2__Alphaproteobacteria;D_3__Rhodobacterales;D_4__Rhodobacteraceae | 0 | 0 | 0 | 0 | 0 | 0 | 0 | 0 | 0 | 0 | 0 | 0 | 0 | 0 | 0 | 0 | 0 | 0 | 0 | 0 | 0 | 15 | 0 | 0 | 0 | 0 | 0 | 0 | 0 | 0 | 0 | 0 | 0 | 0 | 0 | 9 | 24 | 0.002016 |
| D_0__Bacteria;D_1__Proteobacteria;D_2__Gammaproteobacteria;D_3__Vibrionales;D_4__Vibrionaceae | 0 | 0 | 0 | 0 | 0 | 0 | 0 | 0 | 0 | 0 | 0 | 0 | 0 | 0 | 0 | 0 | 0 | 0 | 0 | 0 | 0 | 0 | 0 | 0 | 0 | 0 | 0 | 0 | 0 | 0 | 0 | 0 | 0 | 13 | 11 | 0 | 24 | 0.002016 |
| D_0__Bacteria;D_1__Firmicutes;D_2__Clostridia;D_3__Clostridiales;D_4__Family XI | 0 | 0 | 0 | 0 | 0 | 0 | 0 | 0 | 0 | 0 | 0 | 0 | 0 | 0 | 0 | 0 | 0 | 0 | 0 | 0 | 0 | 10 | 0 | 0 | 3 | 0 | 0 | 0 | 0 | 0 | 0 | 0 | 0 | 0 | 0 | 0 | 13 | 0.001092 |
| D_0__Bacteria;D_1__Patescibacteria;D_2__Saccharimonadia;D_3__Saccharimonadales;D_4__uncultured bacterium | 0 | 0 | 0 | 0 | 0 | 0 | 0 | 0 | 0 | 0 | 0 | 0 | 0 | 0 | 0 | 0 | 0 | 0 | 0 | 0 | 0 | 0 | 0 | 0 | 0 | 0 | 0 | 0 | 0 | 0 | 0 | 0 | 0 | 6 | 0 | 4 | 10 | 0.00084 |
| D_0__Bacteria;D_1__Firmicutes;D_2__Bacilli;D_3__Lactobacillales;D_4__Aerococcaceae | 0 | 4 | 0 | 0 | 0 | 0 | 0 | 0 | 0 | 0 | 0 | 0 | 0 | 0 | 0 | 0 | 0 | 0 | 0 | 0 | 0 | 0 | 0 | 0 | 0 | 0 | 0 | 0 | 0 | 0 | 0 | 5 | 0 | 0 | 0 | 0 | 9 | 0.000756 |
| D_0__Bacteria;D_1__Bacteroidetes;D_2__Bacteroidia;D_3__Flavobacteriales;D_4__Flavobacteriaceae | 0 | 2 | 0 | 0 | 0 | 0 | 0 | 0 | 0 | 0 | 0 | 0 | 0 | 0 | 0 | 0 | 0 | 0 | 0 | 0 | 0 | 0 | 0 | 0 | 0 | 0 | 0 | 0 | 0 | 0 | 0 | 0 | 0 | 0 | 0 | 0 | 2 | 0.000168 |

**Supplementary S6**

*T. molitor*

| | | | | | | | |
|---|---|---|---|---|---|---|---|
| 100%ofsamples | | | | | | | |
| Feature ID | 2% | 9% | 25% | 50% | 75% | 91% | 98% |
| D_0__Bacteria;D_1__Tenericutes;D_2__Mollicutes;D_3__Entomoplasmatales;D_4__Spiroplasmataceae;D_5__Spiroplasma;D_6__uncultured Spiroplasma sp. | 108.2 | 273.4 | 651 | 23608 | 38182 | 38720.24 | 38955.72 |
| D_0__Bacteria;D_1__Proteobacteria;D_2__Gammaproteobacteria;D_3__Enterobacteriales;D_4__Enterobacteriaceae;__;__ | 373.44 | 420.48 | 528 | 13517 | 21499 | 22918.52 | 23539.56 |
| D_0__Bacteria;D_1__Firmicutes;D_2__Bacilli;D_3__Lactobacillales;D_4__Enterococcaceae;D_5__Enterococcus;__ | 29.44 | 97.48 | 253 | 1607 | 2968 | 3427.52 | 3628.56 |
| D_0__Bacteria;D_1__Firmicutes;D_2__Bacilli;D_3__Bacillales;D_4__Staphylococcaceae;D_5__Staphylococcus;__ | 42.44 | 117.48 | 289 | 1209 | 1811 | 2555.96 | 2881.88 |
| D_0__Bacteria;D_1__Proteobacteria;D_2__Gammaproteobacteria;D_3__Enterobacteriales;D_4__Enterobacteriaceae;D_5__Enterobacter;__ | 9.92 | 30.64 | 78 | 565 | 621 | 626.76 | 629.28 |
| D_0__Bacteria;D_1__Firmicutes;D_2__Bacilli;D_3__Lactobacillales;D_4__Streptococcaceae;D_5__Lactococcus;D_6__uncultured bacterium | 2.24 | 3.08 | 5 | 443 | 465 | 565.48 | 609.44 |
| D_0__Bacteria;D_1__Proteobacteria;D_2__Gammaproteobacteria;D_3__Pseudomonadales;D_4__Pseudomonadaceae;D_5__Pseudomonas;__ | 73.28 | 112.76 | 203 | 330 | 339 | 395.32 | 419.96 |
| D_0__Bacteria;D_1__Firmicutes;D_2__Bacilli;D_3__Bacillales;D_4__Bacillaceae;D_5__Bacillus;__ | 72.32 | 80.44 | 99 | 177 | 181 | 638.6 | 838.8 |
| D_0__Bacteria;D_1__Proteobacteria;D_2__Gammaproteobacteria;D_3__Enterobacteriales;D_4__Enterobacteriaceae;D_5__Serratia;__ | 5.24 | 20.08 | 54 | 71 | 183 | 327 | 390 |
| D_0__Bacteria;D_1__Proteobacteria;D_2__Gammaproteobacteria;D_3__Enterobacteriales;D_4__Enterobacteriaceae;D_5__Pantoea;__ | 11.08 | 11.36 | 12 | 43 | 70 | 89.2 | 97.6 |
| 70%ofsamples | | | | | | | |
| Feature ID | 2% | 9% | 25% | 50% | 75% | 91% | 98% |
| D_0__Bacteria;D_1__Tenericutes;D_2__Mollicutes;D_3__Entomoplasmatales;D_4__Spiroplasmataceae;D_5__Spiroplasma;D_6__uncultured Spiroplasma sp. | 108.2 | 273.4 | 651 | 23608 | 38182 | 38720.24 | 38955.72 |
| D_0__Bacteria;D_1__Proteobacteria;D_2__Gammaproteobacteria;D_3__Enterobacteriales;D_4__Enterobacteriaceae;__;__ | 373.44 | 420.48 | 528 | 13517 | 21499 | 22918.52 | 23539.56 |
| D_0__Bacteria;D_1__Firmicutes;D_2__Bacilli;D_3__Lactobacillales;D_4__Streptococcaceae;D_5__Lactococcus;__ | 0.4 | 1.8 | 5 | 1846 | 1974 | 2781.04 | 3134.12 |
| D_0__Bacteria;D_1__Firmicutes;D_2__Bacilli;D_3__Lactobacillales;D_4__Enterococcaceae;D_5__Enterococcus;__ | 29.44 | 97.48 | 253 | 1607 | 2968 | 3427.52 | 3628.56 |
| D_0__Bacteria;D_1__Firmicutes;D_2__Bacilli;D_3__Lactobacillales;D_4__Lactobacillaceae;D_5__Lactobacillus;D_6__uncultured bacterium | 6.96 | 31.32 | 87 | 1268 | 1938 | 2294.48 | 2450.44 |
| D_0__Bacteria;D_1__Firmicutes;D_2__Bacilli;D_3__Bacillales;D_4__Staphylococcaceae;D_5__Staphylococcus;__ | 42.44 | 117.48 | 289 | 1209 | 1811 | 2555.96 | 2881.88 |
| D_0__Bacteria;D_1__Firmicutes;D_2__Bacilli;D_3__Lactobacillales;D_4__Leuconostocaceae;D_5__Weissella;D_6__Weissella confusa | 0.24 | 1.08 | 3 | 661 | 1114 | 1327.12 | 1420.36 |
| D_0__Bacteria;D_1__Firmicutes;D_2__Bacilli;D_3__Lactobacillales;D_4__Lactobacillaceae;D_5__Lactobacillus;__ | 0.56 | 2.52 | 7 | 626 | 766 | 1038 | 1157 |
| D_0__Bacteria;D_1__Proteobacteria;D_2__Gammaproteobacteria;D_3__Enterobacteriales;D_4__Enterobacteriaceae;D_5__Enterobacter;__ | 9.92 | 30.64 | 78 | 565 | 621 | 626.76 | 629.28 |
| D_0__Bacteria;D_1__Firmicutes;D_2__Bacilli;D_3__Bacillales;D_4__Staphylococcaceae;D_5__Staphylococcus;D_6__uncultured bacterium | 12 | 54 | 150 | 464 | 751 | 1063.32 | 1199.96 |
| D_0__Bacteria;D_1__Firmicutes;D_2__Bacilli;D_3__Lactobacillales;D_4__Streptococcaceae;D_5__Lactococcus;D_6__uncultured bacterium | 2.24 | 3.08 | 5 | 443 | 465 | 565.48 | 609.44 |
| D_0__Bacteria;D_1__Firmicutes;D_2__Bacilli;D_3__Lactobacillales;D_4__Lactobacillaceae;D_5__Pediococcus;D_6__Pediococcus pentosaceus | 0.08 | 0.36 | 1 | 389 | 480 | 613.12 | 671.36 |
| D_0__Bacteria;D_1__Proteobacteria;D_2__Gammaproteobacteria;D_3__Pseudomonadales;D_4__Pseudomonadaceae;D_5__Pseudomonas;__ | 73.28 | 112.76 | 203 | 330 | 339 | 395.32 | 419.96 |
| D_0__Bacteria;D_1__Proteobacteria;D_2__Gammaproteobacteria;D_3__Pseudomonadales;D_4__Pseudomonadaceae;D_5__Pseudomonas;D_6__Pseudomonas aeruginosa | 7.52 | 33.84 | 94 | 183 | 186 | 339.6 | 406.8 |
| D_0__Bacteria;D_1__Firmicutes;D_2__Bacilli;D_3__Bacillales;D_4__Bacillaceae;D_5__Bacillus;__ | 72.32 | 80.44 | 99 | 177 | 181 | 638.6 | 838.8 |
| D_0__Bacteria;D_1__Proteobacteria;D_2__Gammaproteobacteria;D_3__Pseudomonadales;D_4__Pseudomonadaceae;D_5__Pseudomonas;D_6__uncultured bacterium | 1.76 | 7.92 | 22 | 75 | 118 | 182.64 | 210.92 |
| D_0__Bacteria;D_1__Proteobacteria;D_2__Gammaproteobacteria;D_3__Enterobacteriales;D_4__Enterobacteriaceae;D_5__Serratia;__ | 5.24 | 20.08 | 54 | 71 | 183 | 327 | 390 |
| Unassigned;__;__;__;__;__;__ | 3.6 | 16.2 | 45 | 52 | 71 | 172.12 | 216.36 |
| D_0__Bacteria;D_1__Proteobacteria;D_2__Gammaproteobacteria;D_3__Enterobacteriales;D_4__Enterobacteriaceae;D_5__Pantoea;__ | 11.08 | 11.36 | 12 | 43 | 70 | 89.2 | 97.6 |
| D_0__Bacteria;D_1__Actinobacteria;D_2__Actinobacteria;D_3__Micrococcales;D_4__Micrococcaceae;D_5__Kocuria;__ | 0.56 | 2.52 | 7 | 41 | 71 | 90.84 | 99.52 |
| D_0__Bacteria;D_1__Firmicutes;D_2__Bacilli;D_3__Lactobacillales;D_4__Leuconostocaceae;D_5__Leuconostoc;D_6__Leuconostoc pseudomesenteroides | 0.24 | 1.08 | 3 | 28 | 33 | 39.4 | 42.2 |

#### *A. diaperinus*

| 70%and100%ofsamples | | | | | | | |
|---|---|---|---|---|---|---|---|
| Feature ID | 2% | 9% | 25% | 50% | 75% | 91% | 98% |
| D_0__Bacteria;D_1__Proteobacteria;D_2__Gammaproteobacteria;D_3__Enterobacteriales;D_4__Enterobacteriaceae;D_5__Enterobacter;__ | 3451.48 | 3547.66 | 3767.5 | 4111 | 50509 | 80203.72 | 93195.16 |
| D_0__Bacteria;D_1__Proteobacteria;D_2__Gammaproteobacteria;D_3__Enterobacteriales;D_4__Enterobacteriaceae;__;__ | 799.08 | 1089.86 | 1754.5 | 2793 | 2841.5 | 2872.54 | 2886.12 |
| D_0__Bacteria;D_1__Firmicutes;D_2__Bacilli;D_3__Lactobacillales;D_4__Enterococcaceae;D_5__Enterococcus;__ | 356.6 | 376.2 | 421 | 491 | 1392 | 1968.64 | 2220.92 |
| D_0__Bacteria;D_1__Firmicutes;D_2__Bacilli;D_3__Bacillales;D_4__Staphylococcaceae;D_5__Staphylococcus;__ | 232.96 | 267.82 | 347.5 | 472 | 903.5 | 1179.66 | 1300.48 |
| D_0__Bacteria;D_1__Firmicutes;D_2__Bacilli;D_3__Lactobacillales;D_4__Streptococcaceae;D_5__Lactococcus;D_6__uncultured bacterium | 131.92 | 156.14 | 211.5 | 298 | 1386.5 | 2083.14 | 2387.92 |
| D_0__Bacteria;D_1__Firmicutes;D_2__Bacilli;D_3__Lactobacillales;D_4__Streptococcaceae;D_5__Lactococcus;__ | 157.24 | 165.08 | 183 | 211 | 1049 | 1585.32 | 1819.96 |
| D_0__Bacteria;D_1__Firmicutes;D_2__Bacilli;D_3__Lactobacillales;D_4__Enterococcaceae;D_5__Enterococcus;D_6__Enterococcus faecalis | 15.56 | 17.52 | 22 | 29 | 335.5 | 531.66 | 617.48 |
| D_0__Bacteria;D_1__Firmicutes;D_2__Bacilli;D_3__Bacillales;D_4__Listeriaceae;D_5__Listeria;__ | 15.24 | 16.08 | 18 | 21 | 244.5 | 387.54 | 450.12 |
| D_0__Bacteria;D_1__Actinobacteria;D_2__Actinobacteria;D_3__Micrococcales;D_4__Brevibacteriaceae;D_5__Brevibacterium;__ | 13.08 | 13.36 | 14 | 15 | 140.5 | 220.82 | 255.96 |
| D_0__Bacteria;D_1__Actinobacteria;D_2__Actinobacteria;D_3__Corynebacteriales;D_4__Corynebacteriaceae;D_5__Corynebacterium 1;__ | 10.16 | 10.72 | 12 | 14 | 208 | 332.16 | 386.48 |
| D_0__Bacteria;D_1__Actinobacteria;D_2__Actinobacteria;D_3__Micrococcales;D_4__Dermabacteraceae;D_5__Brachybacterium;__ | 6.32 | 7.44 | 10 | 14 | 100.5 | 155.86 | 180.08 |
| D_0__Bacteria;D_1__Firmicutes;D_2__Bacilli;D_3__Bacillales;D_4__Bacillaceae;D_5__Bacillus;__ | 4.04 | 4.18 | 4.5 | 5 | 5 | 5 | 5 |
| D_0__Bacteria;D_1__Proteobacteria;D_2__Gammaproteobacteria;D_3__Pseudomonadales;D_4__Moraxellaceae;D_5__Acinetobacter;__ | 2.08 | 2.36 | 3 | 4 | 17.5 | 26.14 | 29.92 |
| D_0__Bacteria;D_1__Firmicutes;D_2__Bacilli;D_3__Bacillales;D_4__Bacillaceae;D_5__Bacillus;D_6__Bacillus pumilus | 3 | 3 | 3 | 3 | 4 | 4.64 | 4.92 |

*A. domesticus*

| 100%ofsamples | | | | | | | |
|---|---|---|---|---|---|---|---|
| Feature ID | 2% | 9% | 25% | 50% | 75% | 91% | 98% |
| D_0__Bacteria;D_1__Bacteroidetes;D_2__Bacteroidia;D_3__Bacteroidales;D_4__Tannerellaceae;D_5__Parabacteroides;D_6__uncultured bacterium | 327.68 | 407.06 | 588.5 | 725 | 1343.75 | 2451.11 | 2935.58 |
| D_0__Bacteria;D_1__Bacteroidetes;D_2__Bacteroidia;D_3__Bacteroidales;D_4__Bacteroidaceae;D_5__Bacteroides;__ | 93.32 | 129.44 | 212 | 294 | 850.25 | 1843.37 | 2277.86 |
| D_0__Bacteria;D_1__Proteobacteria;D_2__Gammaproteobacteria;D_3__Enterobacteriales;D_4__Enterobacteriaceae;__;__ | 116.76 | 126.42 | 148.5 | 219 | 349.5 | 486.78 | 546.84 |
| D_0__Bacteria;D_1__Firmicutes;D_2__Clostridia;D_3__Clostridiales;D_4__Lachnospiraceae;__;__ | 89.66 | 102.47 | 131.75 | 149.5 | 184.5 | 246.9 | 274.2 |
| D_0__Bacteria;D_1__Bacteroidetes;D_2__Bacteroidia;D_3__Bacteroidales;D_4__Tannerellaceae;D_5__Parabacteroides;__ | 71.12 | 82.04 | 107 | 139 | 369 | 774.12 | 951.36 |
| D_0__Bacteria;D_1__Bacteroidetes;D_2__Bacteroidia;D_3__Bacteroidales;D_4__Rikenellaceae;D_5__Alistipes;__ | 31.1 | 38.45 | 55.25 | 108.5 | 170.5 | 204.1 | 218.8 |
| D_0__Bacteria;D_1__Firmicutes;D_2__Bacilli;D_3__Lactobacillales;D_4__Enterococcaceae;D_5__Enterococcus;__ | 64.2 | 68.4 | 78 | 90.5 | 463 | 1163.8 | 1470.4 |
| D_0__Bacteria;D_1__Bacteroidetes;D_2__Bacteroidia;D_3__Bacteroidales;D_4__Dysgonomonadaceae;D_5__Dysgonomonas;__ | 25.12 | 36.04 | 61 | 79 | 96 | 119.04 | 129.12 |
| D_0__Bacteria;D_1__Proteobacteria;D_2__Gammaproteobacteria;D_3__Enterobacteriales;D_4__Enterobacteriaceae;D_5__Enterobacter;__ | 19.82 | 29.69 | 52.25 | 71.5 | 103.75 | 151.27 | 172.06 |
| D_0__Bacteria;D_1__Firmicutes;D_2__Clostridia;D_3__Clostridiales;D_4__Ruminococcaceae;D_5__uncultured;__ | 55.24 | 56.08 | 58 | 63.5 | 74.75 | 87.71 | 93.38 |
| D_0__Bacteria;D_1__Firmicutes;D_2__Clostridia;D_3__Clostridiales;D_4__Ruminococcaceae;D_5__GCA-900066225;__ | 19.38 | 24.21 | 35.25 | 56 | 71.25 | 71.73 | 71.94 |
| D_0__Bacteria;D_1__Proteobacteria;D_2__Gammaproteobacteria;D_3__Enterobacteriales;D_4__Enterobacteriaceae;D_5__Klebsiella;__ | 11.46 | 20.07 | 39.75 | 51.5 | 58.25 | 68.33 | 72.74 |
| D_0__Bacteria;D_1__Bacteroidetes;D_2__Bacteroidia;D_3__Bacteroidales;D_4__Rikenellaceae;D_5__Alistipes;D_6__uncultured bacterium | 23.42 | 24.89 | 28.25 | 50.5 | 72.75 | 76.11 | 77.58 |
| D_0__Bacteria;D_1__Proteobacteria;D_2__Gammaproteobacteria;D_3__Pseudomonadales;D_4__Pseudomonadaceae;D_5__Pseudomonas;__ | 43.36 | 44.62 | 47.5 | 49.5 | 75.25 | 123.73 | 144.94 |
| D_0__Bacteria;D_1__Proteobacteria;D_2__Gammaproteobacteria;D_3__Enterobacteriales;D_4__Enterobacteriaceae;D_5__Citrobacter;__ | 11.98 | 18.91 | 34.75 | 47.5 | 59.5 | 73.9 | 80.2 |
| D_0__Bacteria;D_1__Firmicutes;D_2__Clostridia;D_3__Clostridiales;D_4__Ruminococcaceae;D_5__Candidatus Soleaferrea;__ | 25.12 | 25.54 | 26.5 | 35 | 44.5 | 47.38 | 48.64 |
| D_0__Bacteria;D_1__Firmicutes;D_2__Bacilli;D_3__Lactobacillales;D_4__Streptococcaceae;D_5__Lactococcus;D_6__uncultured bacterium | 14.6 | 16.7 | 21.5 | 34.5 | 289 | 757.48 | 962.44 |
| D_0__Bacteria;D_1__Proteobacteria;D_2__Deltaproteobacteria;D_3__Desulfovibrionales;D_4__Desulfovibrionaceae;D_5__Desulfovibrio;__ | 12.3 | 13.35 | 15.75 | 24.5 | 41 | 58.28 | 65.84 |
| D_0__Bacteria;D_1__Firmicutes;D_2__Clostridia;D_3__Clostridiales;D_4__Ruminococcaceae;D_5__Candidatus Soleaferrea;D_6__uncultured bacterium | 2.36 | 3.62 | 6.5 | 16 | 26 | 29.84 | 31.52 |
| D_0__Bacteria;D_1__Firmicutes;D_2__Clostridia;D_3__Clostridiales;D_4__Lachnospiraceae;D_5__uncultured;D_6__uncultured bacterium | 6.24 | 7.08 | 9 | 16 | 31.5 | 49.74 | 57.72 |
| D_0__Bacteria;D_1__Firmicutes;D_2__Clostridia;D_3__Clostridiales;D_4__Family XIII;D_5__Anaerovorax;__ | 5.18 | 5.81 | 7.25 | 14 | 20 | 20 | 20 |
| D_0__Bacteria;D_1__Firmicutes;D_2__Negativicutes;D_3__Selenomonadales;D_4__Acidaminococcaceae;D_5__uncultured;__ | 3.6 | 5.7 | 10.5 | 13 | 13.75 | 15.19 | 15.82 |
| D_0__Bacteria;D_1__Verrucomicrobia;D_2__Verrucomicrobiae;D_3__Verrucomicrobiales;D_4__Akkermansiaceae;D_5__Akkermansia;D_6__uncultured bacterium | 5.3 | 6.35 | 8.75 | 13 | 28 | 51.04 | 61.12 |
| D_0__Bacteria;D_1__Bacteroidetes;D_2__Bacteroidia;D_3__Bacteroidales;D_4__Paludibacteraceae;D_5__Paludibacter;__ | 8.06 | 8.27 | 8.75 | 11 | 15.75 | 21.03 | 23.34 |
| D_0__Bacteria;D_1__Firmicutes;D_2__Erysipelotrichia;D_3__Erysipelotrichales;D_4__Erysipelotrichaceae;D_5__Erysipelatoclostridium;D_6__uncultured bacterium | 5.18 | 5.81 | 7.25 | 10.5 | 13 | 13 | 13 |
| D_0__Bacteria;D_1__Firmicutes;D_2__Clostridia;D_3__Clostridiales;D_4__Lachnospiraceae;D_5__Lachnoclostridium;__ | 5 | 5 | 5 | 8.5 | 13.25 | 15.65 | 16.7 |
| D_0__Bacteria;D_1__Proteobacteria;D_2__Alphaproteobacteria;D_3__Rhodospirillales;D_4__Magnetospirillaceae;__;__ | 2.3 | 3.35 | 5.75 | 7.5 | 8 | 8 | 8 |
| D_0__Bacteria;D_1__Firmicutes;D_2__Clostridia;D_3__Clostridiales;D_4__Lachnospiraceae;D_5__GCA-900066575;D_6__uncultured bacterium | 4.06 | 4.27 | 4.75 | 5 | 7.25 | 11.57 | 13.46 |
| D_0__Bacteria;D_1__Firmicutes;D_2__Erysipelotrichia;D_3__Erysipelotrichales;D_4__Erysipelotrichaceae;D_5__Erysipelatoclostridium;__ | 1.12 | 1.54 | 2.5 | 3 | 7.25 | 15.41 | 18.98 |

| 70%ofsamples | | | | | | | |
|---|---|---|---|---|---|---|---|
| Feature ID | 2% | 9% | 25% | 50% | 75% | 91% | 98% |
| D_0__Bacteria;D_1__Bacteroidetes;D_2__Bacteroidia;D_3__Bacteroidales;D_4__Tannerellaceae;D_5__Parabacteroides;D_6__uncultured bacterium | 327.68 | 407.06 | 588.5 | 725 | 1343.75 | 2451.11 | 2935.58 |
| D_0__Bacteria;D_1__Firmicutes;D_2__Bacilli;D_3__Bacillales;D_4__Bacillaceae;D_5__Bacillus;__ | 1.14 | 5.13 | 14.25 | 355 | 1693.75 | 3619.03 | 4461.34 |
| D_0__Bacteria;D_1__Bacteroidetes;D_2__Bacteroidia;D_3__Bacteroidales;D_4__Bacteroidaceae;D_5__Bacteroides;__ | 93.32 | 129.44 | 212 | 294 | 850.25 | 1843.37 | 2277.86 |
| D_0__Bacteria;D_1__Proteobacteria;D_2__Gammaproteobacteria;D_3__Enterobacteriales;D_4__Enterobacteriaceae;__;__ | 116.76 | 126.42 | 148.5 | 219 | 349.5 | 486.78 | 546.84 |
| D_0__Bacteria;D_1__Firmicutes;D_2__Clostridia;D_3__Clostridiales;D_4__Lachnospiraceae;__;__ | 89.66 | 102.47 | 131.75 | 149.5 | 184.5 | 246.9 | 274.2 |
| D_0__Bacteria;D_1__Bacteroidetes;D_2__Bacteroidia;D_3__Bacteroidales;D_4__Tannerellaceae;D_5__Parabacteroides;__ | 71.12 | 82.04 | 107 | 139 | 369 | 774.12 | 951.36 |
| D_0__Bacteria;D_1__Bacteroidetes;D_2__Bacteroidia;D_3__Bacteroidales;D_4__Rikenellaceae;D_5__Alistipes;__ | 31.1 | 38.45 | 55.25 | 108.5 | 170.5 | 204.1 | 218.8 |
| D_0__Bacteria;D_1__Firmicutes;D_2__Bacilli;D_3__Lactobacillales;D_4__Enterococcaceae;D_5__Enterococcus;__ | 64.2 | 68.4 | 78 | 90.5 | 463 | 1163.8 | 1470.4 |
| D_0__Bacteria;D_1__Bacteroidetes;D_2__Bacteroidia;D_3__Bacteroidales;D_4__Dysgonomonadaceae;D_5__Dysgonomonas;__ | 25.12 | 36.04 | 61 | 79 | 96 | 119.04 | 129.12 |
| D_0__Bacteria;D_1__Proteobacteria;D_2__Gammaproteobacteria;D_3__Enterobacteriales;D_4__Enterobacteriaceae;D_5__Enterobacter;__ | 19.82 | 29.69 | 52.25 | 71.5 | 103.75 | 151.27 | 172.06 |
| D_0__Bacteria;D_1__Firmicutes;D_2__Clostridia;D_3__Clostridiales;D_4__Ruminococcaceae;D_5__uncultured;__ | 55.24 | 56.08 | 58 | 63.5 | 74.75 | 87.71 | 93.38 |
| D_0__Bacteria;D_1__Firmicutes;D_2__Clostridia;D_3__Clostridiales;D_4__Ruminococcaceae;D_5__GCA-900066225;__ | 19.38 | 24.21 | 35.25 | 56 | 71.25 | 71.73 | 71.94 |
| D_0__Bacteria;D_1__Proteobacteria;D_2__Gammaproteobacteria;D_3__Enterobacteriales;D_4__Enterobacteriaceae;D_5__Klebsiella;__ | 11.46 | 20.07 | 39.75 | 51.5 | 58.25 | 68.33 | 72.74 |
| D_0__Bacteria;D_1__Bacteroidetes;D_2__Bacteroidia;D_3__Bacteroidales;D_4__Rikenellaceae;D_5__Alistipes;D_6__uncultured bacterium | 23.42 | 24.89 | 28.25 | 50.5 | 72.75 | 76.11 | 77.58 |
| D_0__Bacteria;D_1__Proteobacteria;D_2__Gammaproteobacteria;D_3__Pseudomonadales;D_4__Pseudomonadaceae;D_5__Pseudomonas;__ | 43.36 | 44.62 | 47.5 | 49.5 | 75.25 | 123.73 | 144.94 |
| D_0__Bacteria;D_1__Proteobacteria;D_2__Gammaproteobacteria;D_3__Enterobacteriales;D_4__Enterobacteriaceae;D_5__Citrobacter;__ | 11.98 | 18.91 | 34.75 | 47.5 | 59.5 | 73.9 | 80.2 |
| D_0__Bacteria;D_1__Firmicutes;D_2__Clostridia;D_3__Clostridiales;D_4__Ruminococcaceae;D_5__Candidatus Soleaferrea;__ | 25.12 | 25.54 | 26.5 | 35 | 44.5 | 47.38 | 48.64 |

| | | | | | | | |
|---|---|---|---|---|---|---|---|
| D_0__Bacteria;D_1__Firmicutes;D_2__Bacilli;D_3__Lactobacillales;D_4__Streptococcaceae;D_5__Lactococcus;D_6__uncultured bacterium | 14.6 | 16.7 | 21.5 | 34.5 | 289 | 757.48 | 962.44 |
| D_0__Bacteria;D_1__Proteobacteria;D_2__Deltaproteobacteria;D_3__Desulfovibrionales;D_4__Desulfovibrionaceae;D_5__Desulfovibrio;__ | 12.3 | 13.35 | 15.75 | 24.5 | 41 | 58.28 | 65.84 |
| D_0__Bacteria;D_1__Firmicutes;D_2__Clostridia;D_3__Clostridiales;D_4__Lachnospiraceae;D_5__Tyzzerella 3;__ | 0.84 | 3.78 | 10.5 | 20 | 26.75 | 28.19 | 28.82 |
| D_0__Bacteria;D_1__Firmicutes;D_2__Clostridia;D_3__Clostridiales;D_4__Ruminococcaceae;D_5__Candidatus Soleaferrea;D_6__uncultured bacterium | 2.36 | 3.62 | 6.5 | 16 | 26 | 29.84 | 31.52 |
| D_0__Bacteria;D_1__Firmicutes;D_2__Clostridia;D_3__Clostridiales;D_4__Lachnospiraceae;D_5__uncultured;D_6__uncultured bacterium | 6.24 | 7.08 | 9 | 16 | 31.5 | 49.74 | 57.72 |
| D_0__Bacteria;D_1__Firmicutes;D_2__Clostridia;D_3__Clostridiales;D_4__Family XIII;D_5__Anaerovorax;__ | 5.18 | 5.81 | 7.25 | 14 | 20 | 20 | 20 |
| D_0__Bacteria;D_1__Verrucomicrobia;D_2__Verrucomicrobiae;D_3__Verrucomicrobiales;D_4__Akkermansiaceae;D_5__Akkermansia;D_6__uncultured bacterium | 5.3 | 6.35 | 8.75 | 13 | 28 | 51.04 | 61.12 |
| D_0__Bacteria;D_1__Firmicutes;D_2__Negativicutes;D_3__Selenomonadales;D_4__Acidaminococcaceae;D_5__uncultured;__ | 3.6 | 5.7 | 10.5 | 13 | 13.75 | 15.19 | 15.82 |
| D_0__Bacteria;D_1__Bacteroidetes;D_2__Bacteroidia;D_3__Flavobacteriales;D_4__Weeksellaceae;D_5__Apibacter;__ | 0.54 | 2.43 | 6.75 | 12 | 20 | 29.6 | 33.8 |
| D_0__Bacteria;D_1__Bacteroidetes;D_2__Bacteroidia;D_3__Bacteroidales;D_4__Paludibacteraceae;D_5__Paludibacter;__ | 8.06 | 8.27 | 8.75 | 11 | 15.75 | 21.03 | 23.34 |
| D_0__Bacteria;D_1__Firmicutes;D_2__Erysipelotrichia;D_3__Erysipelotrichales;D_4__Erysipelotrichaceae;D_5__Erysipelatoclostridium;D_6__uncultured bacterium | 5.18 | 5.81 | 7.25 | 10.5 | 13 | 13 | 13 |
| D_0__Bacteria;D_1__Firmicutes;D_2__Clostridia;D_3__Clostridiales;D_4__Lachnospiraceae;D_5__Lachnoclostridium;__ | 5 | 5 | 5 | 8.5 | 13.25 | 15.65 | 16.7 |
| D_0__Bacteria;D_1__Firmicutes;D_2__Clostridia;D_3__Clostridiales;D_4__Ruminococcaceae;D_5__Butyricicoccus;__ | 0.48 | 2.16 | 6 | 8 | 14.5 | 26.98 | 32.44 |
| D_0__Bacteria;D_1__Firmicutes;D_2__Clostridia;D_3__Clostridiales;D_4__Lachnospiraceae;D_5__Blautia;__ | 0.36 | 1.62 | 4.5 | 8 | 13.25 | 19.49 | 22.22 |
| D_0__Bacteria;D_1__Bacteroidetes;D_2__Bacteroidia;D_3__Bacteroidales;D_4__Dysgonomonadaceae;D_5__uncultured;__ | 0.3 | 1.35 | 3.75 | 7.5 | 11.25 | 13.65 | 14.7 |
| D_0__Bacteria;D_1__Proteobacteria;D_2__Alphaproteobacteria;D_3__Rhodospirillales;D_4__Magnetospirillaceae;__;__ | 2.3 | 3.35 | 5.75 | 7.5 | 8 | 8 | 8 |
| D_0__Bacteria;D_1__Proteobacteria;D_2__Gammaproteobacteria;D_3__Enterobacteriales;D_4__Enterobacteriaceae;D_5__Dickeya;__ | 0.18 | 0.81 | 2.25 | 6 | 86.75 | 236.03 | 301.34 |
| D_0__Bacteria;D_1__Proteobacteria;D_2__Gammaproteobacteria;D_3__Betaproteobacteriales;D_4__Neisseriaceae;__;__ | 0.06 | 0.27 | 0.75 | 5.5 | 10.5 | 11.46 | 11.88 |
| D_0__Bacteria;D_1__Firmicutes;D_2__Clostridia;D_3__Clostridiales;D_4__Lachnospiraceae;D_5__GCA-900066575;D_6__uncultured bacterium | 4.06 | 4.27 | 4.75 | 5 | 7.25 | 11.57 | 13.46 |
| D_0__Bacteria;D_1__Firmicutes;D_2__Clostridia;D_3__Clostridiales;D_4__Lachnospiraceae;D_5__Tyzzerella;__ | 0.12 | 0.54 | 1.5 | 5 | 8.5 | 9.46 | 9.88 |
| D_0__Bacteria;D_1__Fusobacteria;D_2__Fusobacteriia;D_3__Fusobacteriales;D_4__Fusobacteriaceae;D_5__Fusobacterium;__ | 0.12 | 0.54 | 1.5 | 4 | 9.75 | 16.95 | 20.1 |
| D_0__Bacteria;D_1__Proteobacteria;D_2__Deltaproteobacteria;D_3__Desulfovibrionales;D_4__Desulfovibrionaceae;D_5__uncultured;D_6__uncultured bacterium | 0.18 | 0.81 | 2.25 | 4 | 8.75 | 15.95 | 19.1 |
| D_0__Bacteria;D_1__Firmicutes;D_2__Bacilli;D_3__Lactobacillales;D_4__Leuconostocaceae;D_5__Weissella;D_6__Weissella confusa | 0.12 | 0.54 | 1.5 | 3.5 | 50.25 | 137.13 | 175.14 |
| D_0__Bacteria;D_1__Firmicutes;D_2__Erysipelotrichia;D_3__Erysipelotrichales;D_4__Erysipelotrichaceae;D_5__Erysipelatoclostridium;__ | 1.12 | 1.54 | 2.5 | 3 | 7.25 | 15.41 | 18.98 |
| D_0__Bacteria;D_1__Firmicutes;D_2__Clostridia;D_3__Clostridiales;D_4__Lachnospiraceae;D_5__Blautia;D_6__uncultured bacterium | 0.18 | 0.81 | 2.25 | 3 | 5.75 | 11.03 | 13.34 |
| D_0__Bacteria;D_1__Firmicutes;D_2__Clostridia;D_3__Clostridiales;D_4__Christensenellaceae;D_5__Christensenellaceae R-7 group;D_6__uncultured bacterium | 0.12 | 0.54 | 1.5 | 2.5 | 4.75 | 8.11 | 9.58 |
| D_0__Bacteria;D_1__Firmicutes;D_2__Clostridia;D_3__Clostridiales;D_4__Clostridiales vadinBB60 group;D_5__uncultured bacterium;D_6__uncultured bacterium | 0.06 | 0.27 | 0.75 | 2 | 33.75 | 92.79 | 118.62 |

Venn results

| Names | total | genera |
|---|---|---|
| Acheta ∩ Alphitobius ∩ Tenebrio | 4 | Enterobacteriaceae_genus |
| | | Lactococcus |
| | | Enterobacter |
| | | Enterococcus |
| Alphitobius ∩ Tenebrio | 2 | Staphylococcus |
| | | Bacillus |
| Acheta ∩ Tenebric | 1 | Pseudomonas |
| Tenebrio | 3 | Spiroplasma |
| | | Pantoea |
| | | Serratia |
| Alphitobius | 5 | Brevibacterium |
| | | Acinetobacter |
| | | Brachybacterium |
| | | Listeria |
| | | Corynebacterium 1 |
| Acheta | 20 | Paludibacter |
| | | Candidatus_Soleaferrea |
| | | Magnetospirillaceae_genus |
| | | Bacteroides |
| | | Parabacteroides |
| | | Dysgonomonas |
| | | Akkermansia |
| | | Alistipes |
| | | Acidaminococcaceae_uncultured |
| | | Erysipelatoclostridium |
| | | Lachnoclostridium |
| | | Klebsiella |
| | | Lachnospiraceae_GCA-900066575 |
| | | Ruminococcaceae_uncultured |
| | | Ruminococcaceae_GCA-900066225 |
| | | Lachnospiraceae_uncultured |
| | | Citrobacter |
| | | Lachnospiraceae_genus |
| | | Desulfovibrio |
| | | Anaerovorax |
